# Changes in ambient temperature are the prevailing cue in determining *Brachypodium distachyon* diurnal gene regulation

**DOI:** 10.1101/762021

**Authors:** Kirk J-M. MacKinnon, Benjamin J. Cole, Chang Yu, Joshua H. Coomey, Nolan T. Hartwick, Marie-Stanislas Remigereau, Tomás Duffy, Todd P. Michael, Steve A. Kay, Samuel P. Hazen

## Abstract

- Plants are continuously exposed to diurnal fluctuations in light and temperature, and spontaneous changes in their physical or biotic environment. The circadian clock coordinates regulation of gene expression with a 24-hour period, enabling the anticipation of these events.
- We used RNA sequencing to characterize the *Brachypodium distachyon* transcriptome under light and temperature cycles, as well as under constant conditions.
- Approximately 3% of the transcriptome was regulated by the circadian clock, a smaller proportion reported in most other species. For most transcripts that were rhythmic under all conditions, including many known clock genes, the period of gene expression lengthened from 24 to 27 h in the absence of external cues. To functionally characterize the cyclic transcriptome in *B. distachyon*, we used Gene Ontology enrichment analysis, and found several terms significantly associated with peak expression at particular times of the day. Furthermore we identified sequence motifs enriched in the promoters of similarly-phased genes, some potentially associated with transcription factors.
- When considering the overlap in rhythmic gene expression and specific pathway behavior, thermocycles was the prevailing cue that controlled diurnal gene regulation. Taken together, our characterization of the rhythmic *B. distachyon* transcriptome represents a foundational resource with implications in other grass species.

## INTRODUCTION

The external environment that plants experience is acutely variable within a single day. Changes in physiological behavior can reflect a direct and immediate response to stimuli, or an anticipated response for a predictable change in the environment. Both types of responses are adaptive, but there is a distinct advantage to anticipating recurring changes (Green *et al*., 2002; Michael *et al*., 2003; Dodd *et al*., 2005). An endogenous timekeeper, known as the circadian clock, provides the capacity to synchronize behavior with the environment, as well as the means to time seasonal behavior. Plant circadian clock oscillations are created by a set of interconnected transcriptional feedback loops that drive morning- and evening-specific outputs (Nohales & Kay, 2016). A vast majority of the mechanisms that create daily rhythms were resolved through the study of *Arabidopsis thaliana*, with few if any unique mechanisms identified in other eudicots or monocots. Morning expressed Myb transcription factors, CIRCADIAN CLOCK ASSOCIATED 1 (CCA1) and LONG ELONGATED HYPOCOTYL (LHY), form the morning loop and bind to the evening element sequence in the *cis*-regulatory regions of the *PSEUDO RESPONSE REGULATOR* (*PRR*) gene family, including *TIMING OF CAB EXPRESSION 1* (*TOC1*), and repress their expression (Harmer *et al*., 2000; Arsovski *et al*., 2015; Nagel *et al*., 2015; Kamioka *et al*., 2016). CCA1 and LHY also repress their own expression in addition to genes encoding *EARLY FLOWERING 3* (*ELF3*), *ELF4*, and *LUX ARRYTHMO* (*LUX*), which make up the evening complex (Nusinow *et al*., 2011). The PRRs in turn function as repressors of the morning loop and the evening complex, as well as within the PRR family (Pokhilko *et al*., 2012; Gendron *et al*., 2012; Huang *et al*., 2012; Liu *et al*., 2016). Thus, the morning loop, the evening complex, and the PRRs function primarily as repressors of each other (Pokhilko *et al*., 2012). Activation within the clock occurs by removing this repression, either indirectly through light-stimulated protein degradation or transcriptional repression of the repressing clock genes, or directly through transcriptional activation.

While the core circadian oscillator can intrinsically function for many days, exogenous inputs, e.g. light, humidity, or temperature, are necessary to maintain clock synchronicity (Nohales & Kay, 2016). Light is perceived and transmitted by multiple distinct families of photoreceptors, including phytochromes (PHY), cryptochromes (CRY), ZTL/FKF1/LKP2 proteins (ZEITLUPE, FLAVIN-BINDING KELCH REPEAT F-BOX, and LOV KELCH PROTEIN 2), phototropins, and the ultraviolet light receptor, UVR8 (Zoltowski & Imaizumi, 2014; Franklin *et al*., 2014; Casal & Qüesta, 2018). Light-dependent photoreceptor activity, in turn, influences the function of core clock proteins. In the dark, ZTL targets TOC1, PRR5, and CCA1 HIKING EXPEDITION (CHE) proteins for degradation, releasing the repression of the morning-expressed genes, CCA1 and LHY (Más *et al*., 2003; Lee *et al*., 2018). Both cryptochrome and phytochrome activities have been implicated in controlling the morning loop function of PRR7 and PRR9 (Farré *et al*., 2005).

Phytochromes can also play a key role in the function of evening complex proteins, with PhyA signaling inducing ELF4 expression, and PhyB signaling modulating PHYTOCHROME INTERACTING FACTOR 4 (PIF4) through ELF3 function (Li *et al*., 2011; Nusinow *et al*., 2011; Kolmos *et al*., 2011). PhyB also interacts with CCA1, LHY, GI, TOC1, and ELF3 proteins (Liu *et al*., 2001; Yeom *et al*., 2014). ELF3 represses growth, both directly, through inactivation of PIF4 protein, and in a clock dependent manner, transcriptionally repressing *PIF4* as a component of the evening complex (Nieto *et al*., 2015). More generally, light influences the expression of numerous genes, namely *CCA1*, *LHY*, *GIGANTEA* (*GI*), the *NIGHT LIGHT-INDUCIBLE AND CLOCK-REGULATED* (*LNK*) and *LIGHT-REGULATED WD* (*LWD*)(Nohales & Kay, 2016). Light also causes daytime expression of the LHY-like Myb transcription factor *REVEILLE* 8 (*RVE8*) and the LNKs, which activate several evening-expressed clock genes (Rawat *et al*., 2011; Farinas & Mas, 2011; Hsu *et al*., 2013; Rugnone *et al*., 2013; Xie *et al*., 2014). Recently, GI was shown to influence light signaling via direct functional interaction with PIF proteins, further tightening regulatory control of light on clock-mediated transcription (Nohales *et al*., 2019). Thus, the integration of light signals is mediated through several points in the circadian clock, namely through the activation of gene expression, degradation of proteins, or PhyB light- and temperature-dependent function. These light-responsive events are necessary to time daily rhythms of gene expression.

Temperature serves as an entrainment cue that has independent and overlapping effects with light input into the clock. The function of phytochromes is temperature-sensitive; thus, they are thermosensors as well as photoreceptors (Franklin *et al*., 2014). Protein reversion from the active to inactive state, which occurs in the dark, is accelerated by an increase in temperature (Jung *et al*., 2016; Legris *et al*., 2016). Interestingly, PhyB protein interacts with the evening-complex protein ELF3, and *elf3* mutants are incapable of thermocycle entrainment (Reed *et al*., 2000; Thines & Harmon, 2010). Similarly, evening-complex function is temperature-sensitive; direct repression of *PRR7*, *GI*, and *LUX* expression increases as nighttime ambient temperature decreases (Mizuno *et al*., 2014; Box *et al*., 2015). The *cis*-regulatory region of *LUX* is directly bound and activated by the cold-responsive transcription factor CBF1 (C-REPEAT/DRE BINDING FACTOR, also known as DREB) (Chow *et al*., 2014). Many of the CBF downstream targets, such as the *COR* genes, exhibit diurnal and circadian clock regulated expression, peaking at the end of the day (Harmer *et al*., 2000; Michael *et al*., 2008b; Chow *et al*., 2014). The clock in turn gates temperature responsiveness (Lee & Thomashow, 2012). Temperature has also been shown to play a role in post-transcriptional and post-translational regulation of circadian clock components and chromatin state (Kumar & Wigge, 2010; Portolés & Más, 2010; James *et al*., 2012; Seo *et al*., 2012; Choudhary *et al*., 2015; Marshall *et al*., 2016; Gil *et al*., 2017). Thus, numerous mechanisms and molecular outcomes of thermocycles are key to adaptive changes and buffering against spurious responses to temperature fluctuations.

Here, we use RNA-sequencing to profile the abundance of *Brachypodium distachyon* transcripts under diurnal and constant conditions. We observed robust oscillations under photocycles, thermocycles, and both photo- and thermocycles combined. Specific biological functions, such as secondary cell wall biosynthesis and DNA replication, were mostly entrained by only one input. The circadian clock regulated a relatively small proportion of transcripts, many of which exhibited an unusually long period in constant conditions. Together with observed patterns of growth, metabolism, and gene expression demonstrate that grasses may be relatively responsive to thermocycles rather than the circadian clock (Poiré *et al*., 2010; Matos *et al*., 2014; Müller *et al*., 2018).

## MATERIALS AND METHODS

### Plant materials and growth conditions

Bd21 seeds were sterilized as previously described (Weigel & Glazebrook, 2006), stratified at 4 °C for 7 days and sown into 100 mL 1x Murashige and Skoog (MS) medium + 0.8 % agar in Magenta boxes. Seeds were germinated in the dark for 3 days, then transferred to diurnal conditions for 10 days (12 h light, 12 h dark, 28 °C day temperature, 12 °C night temperature). Seedlings were then released into one of four conditions: light:dark hot:cold LDHC (12 h light, 12 °C, 28 h dark, 12 °C), LDHH (12 h light, 12 h dark, continuous 28 °C), LLHC (continuous light, 12 h 28 °C, 12 h 12 °C), or LLHH (continuous light, continuous 28 °C). Light intensity was approximately 50 µmol·m^-2^·sec^-1^. Twelve hours after placing the samples in each condition (the evening of the 10th day, just prior to the anticipated day/night transition), the currently emerging leaf was sampled from at least 3 separate plants, pooled, and frozen in liquid nitrogen. Following this first sample, 13 additional time points (14 in total), were collected, one every 3.5 hours.

### RNA sample preparation and sequencing

Total RNA was extracted as previously described (Handakumbura *et al*., 2018). mRNA was enriched through two rounds of polyA selection using the Dynabeads mRNA Purification kit (ThermoFisher Scientific, Waltham, MA). Illumina stranded, paired-end sequencing libraries were then constructed using the ScriptSeq-v2 RNA-Seq Library Preparation Kit (Illumina, San Diego, CA) following the manufacturer’s instructions. Libraries were sequenced at the USC Epigenome Core Facility (HiSeq 2000; 6 samples on 1 lane; paired-end 100 cycles) or the USC Genome Core Facility (Hiseq 2500; 42 samples on 7 lanes; 6 samples per lane; paired-end, 100 cycles).

### Transcript identification, quantification, and circadian analysis

The processed sequences were aligned to v3.1 of the Bd21 genome using HISAT2 (Kim *et al*., 2015)(**Table S1**). This annotation of the genome contains 34,310 protein-coding loci and 52,972 protein-coding transcripts. Read counts were then normalized using DESeq2 (Love *et al*., 2014). In each time course, approximately 80% of all transcripts were detected, and the distribution of the normalized expression levels were comparable (**Fig. S1**). Unless specified otherwise, all expression quantities reported are at the transcript (as opposed to gene) level, accounting for potential splice variants. After normalization, MetaCycle (Wu *et al*., 2016a) was used to assess for a period that ranged from 21 to 28 h with rhythmic genes called using a *p*-value cutoff of 0.01. Period and phase estimates of genes with a phase of 12 we visually inspected (**Fig. S2**).

### Computational analysis

Computational analysis was performed in R (version 3.6.0) using resources from the Massachusetts Green High Performance Computing Center. Venn diagrams were generated using the Limma package (Ritchie *et al*., 2015). Sinaplots were generated using the ggforce R package (Pedersen, 2019). Remaining plots were created using ggplot2 (Wickham, 2016). Data transformation was accomplished with either the dplyr package (Wickham *et al*., 2019), or the data.table package (Dowle & Srinivasan, 2019).

Coefficient of variance was determined using the raster package (Hijmans, 2019). Tests of statistical significance for period lengths and relative amplitudes were conducted by first performing analysis of variance and then based on the distribution of the data we assessed significance using a pairwise Wilcoxon rank sum test (Haynes, 2013). The transcriptome data were integrated into the Brachypodium eFP browser, (http://bar.utoronto.ca/efp_brachypodium/cgi-bin/efpWeb.cgi?dataSource=Photo_Thermocycle) (Winter *et al*., 2007). The eFP browser enables the visualization of gene expression levels with a pictograph heatmap and the download and visualization of tables and charts for expression values of individual genes. Raw read data were deposited in the European Nucleotide Archive for public access (Accession: <u>PRJEB32498</u>).

### Pathway enrichment analysis

NCBI BLAST and Phytozome (Altschul *et al*., 1990; Goodstein *et al*., 2011) were used to find orthologs for all *B. distachyon* v3.1 genes as the reciprocal best match to *A. thaliana* TAIRv10 protein sequences. Genes that did not significantly match a corresponding gene in *A. thaliana* were discarded from this analysis. *Arabidopsis thaliana* biological process gene ontology (GO) annotations were obtained from http://ge-lab.org/gskb/. The Java implementation of PSEA1.1 (Zhang *et al*., 2016) was used for analysis, with 0-28 for the “domain” parameter, and 10 for the “minimum items” parameter. For Kuiper’s test, the distribution of GO terms was compared with an empirically-determined background distribution. Gene identifiers were submitted to g:Profiler (Raudvere *et al*., 2019) for KEGG and Wiki pathway enrichment analysis.

### Hierarchical cluster analysis

To find groups of genes with similar expression profiles, we first filtered all transcripts to only include those expressed at least once in all four time courses. Transcript expression was then standardized by computing z-scores within each condition. Standardized transcript expression dissimilarity was assessed using Pearson’s correlation and hierarchically clustered according to the complete linkage method of agglomeration. The resulting dendrogram was visually analyzed to determine the ideal number of clusters based on the elbow method. Gene expression was then modeled using the GAM method of smoothing to represent a cluster (Wood, 2004). Period length clustering was performed by first normalizing period values relative to their z-score for transcripts that cycled in all of the four time course conditions. Heatmaps were plotted using Heatmap.2 (Warnes *et al*., 2019). Confidence intervals for period length clusters were tested using ANOVA and post-hoc Tukey’s honest significant difference test. Tests of variance for period distributions by condition were done using Fligner-Killeen test of homogeneity.

### Phylogenetic analysis

To identify *B. distachyon* orthologs of *A. thaliana RVE*s, *LWD*s, and *LNKs*, we used a BLAST e-value ≤ e-60 as the criterion for quickly identifying the immediate homologs for each clade, and larger e-values to extract less related, but phylogenetically relevant sequences. We identified homologs of each clade using *A. thaliana* TAIR10, *B. distachyon* Phytozome v3.1, *Orya sativa* v7.0_JGI, *Populus trichocarpa* Phytozome v3.0, *Setaria viridis* Phytozome v1.1, and *Solanum lycopersicum* ITAG v3.10 reference genomes. To reconstruct phylogenetic relationships between orthologs, we first aligned protein sequences using MAFFT, with the G-INS-I model (Katoh *et al*., 2002) and then used the neighbor-joining method for tree construction with 1000 bootstrap samples. The proteins sequences used are provided in **Table S2**.

### Cis-regulatory sequence analysis

*Cis*-regulatory sequence analysis of cycling transcripts was performed by first grouping transcripts based on phase of gene expression in each time course and then selecting putative regulatory sequence up to 1000 bp upstream of the transcriptional start site. HOMER v4.10 (Heinz *et al*., 2010) was used to compute enrichment scores for transcription factor binding motifs previously identified using DNA-affinity-purified sequencing (DAP-seq, (O’Malley *et al*., 2016)) among each group of cycling transcripts. Motif enrichment was calculated against the hypergeometric distribution; the significance threshold was set to *p* < 0.05. Similar motifs were determined using the compareMotifs.pl function of HOMER against the global list of known motifs with the default threshold cutoff of 0.6. ELEMENT analysis was carried out as previously described with modifications (Mockler *et al*., 2007; Michael *et al*., 2008b; Filichkin *et al*., 2011; Wai *et al*., 2019). Briefly, The *B. distachyon* promoters (500 bp) were parsed using the Bd v3.1 gene models and grouped by peak time of expression into phase bins (0 to 28 hrs).

## RESULTS

### Widespread diurnal regulation of the *Brachypodium distachyon* transcriptome

In order to fully explore the rhythmic nature of the *B. distachyon* transcriptome, accession Bd21 seeds were entrained in photocycles and thermocycles for 10 d after germination. Growth conditions were then shifted to one of three conditions: photocycles only (light:dark hot:hot, LDHH), thermocycles only (light:light hot:cold, LLHC), or constant conditions (light:light hot:hot, LLHH). A fourth set of samples remained in photocycles and thermocycles (light:dark hot:cold, LDHC). After 12 h of exposure to their new conditions, the emerging leaves of plants were sampled every 3.5 h for a total of 45.5 h, resulting in a collection of 14 time-resolved samples across four conditions. Transcript abundances were then quantified from each of these samples using RNA-seq.

Expression patterns of cyclically expressed genes can be broadly summarized by three parameters: phase (the time of peak expression), period (the average time between peaks), and amplitude (half of the gene expression difference between peak and trough). For each expressed transcript (47,036), we estimated the periodicity and phase under each condition using MetaCycle, which integrates three distinct methods: ARSER, JTK.Cycle, and Lomb-Scargle algorithms, combined using Fisher’s method (Yang & Su, 2010; Hughes *et al*., 2010; Wu *et al*., 2016a). The bimodal *p*-value distribution was very similar across growth conditions (**Fig. S3**), enabling us to use a fixed *p*-value cutoff (*p* < 0.01) to classify transcripts as rhythmic without bias towards any specific condition. Based on this analysis, we observed 16,916 transcripts with rhythmic expression in at least one condition, accounting for nearly 36% of the measured transcriptome. The largest number of cycling transcripts, 10,027 (21.3%), was observed in LDHC (**Fig. 1**). The same number of transcripts cycled under photocycles or thermocycles alone (7,391; 15.7%) and the smallest number of rhythmic transcripts was measured under constant conditions (1,699; 3.6%). Among pairs of conditions, LDHC and LLHC conditions were most similar in terms of shared rhythmic transcripts (4,531), while LDHC and LDHH exhibited 30% fewer shared rhythmic transcripts (3,156). We also identified several transcripts that did not cycle under any condition, and might be useful as reference genes for targeted assays ((Hong *et al*., 2008); **Fig. S4**, **Table S3-4**). These results suggest that temperature cycles are more dominant in driving diurnal regulation of the transcriptome.

**Figure 1.**
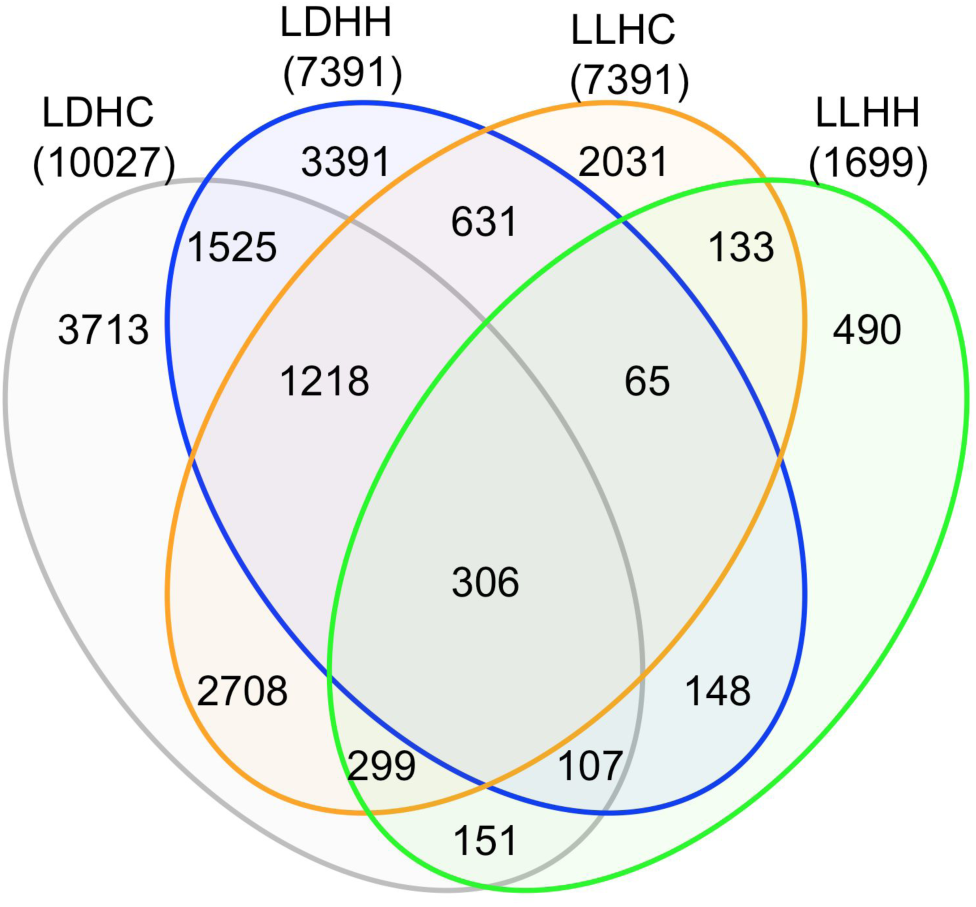
30% of the *Brachypodium distachyon* transcriptome is controlled by photocycles, thermocycles, or the circadian clock. Venn diagram of the number of transcripts determined to be rhythmic by MetaCycle using a p-value cutoff of < 0.01. Each value within the ovals is the count of rhythmic transcripts exclusive to each union. Total numbers of transcripts measured as rhythmic in each condition are in parentheses. LDHC, photo- and thermocycles; LDHH, photocycles and constant temperature; LLHC, thermocycles and constant light; LLHH, constant light and constant warm temperature.

### Circadian clock and photoreceptor genes

Very few circadian clock genes have been functionally characterized in grasses, and none in *B. distachyon*. Therefore, we assembled a list of core circadian clock transcripts based on our analysis of amino acid sequence similarity to *A. thaliana* and by previously described homology and expression behavior in *B. distachyon* (Higgins et al., 2010; Matos et al., 2014; Calixto et al., 2015)(**Fig. S4-6**). To further verify that the *B. distachyon* clock gene orthologs represent true clock genes, we examined whether transcripts from these loci cycle in our transcriptome data. We also examined cryptochrome, phototropin, phytochrome, and ultraviolet photoreceptors and unlike *A. thaliana*, they tended to not be clock regulated (**Fig. S8**, **Table S6**). From this analysis, we observed that all putative clock gene orthologs had at least one rhythmic transcript in at least one of the two-day time courses, with the exception of *BdLWD,* which was starkly arrhythmic (**Fig. 2**, **Table S5**). This observation is consistent with the overall transcriptome behavior where average period was greatest in LLHH.

**Figure 2.**
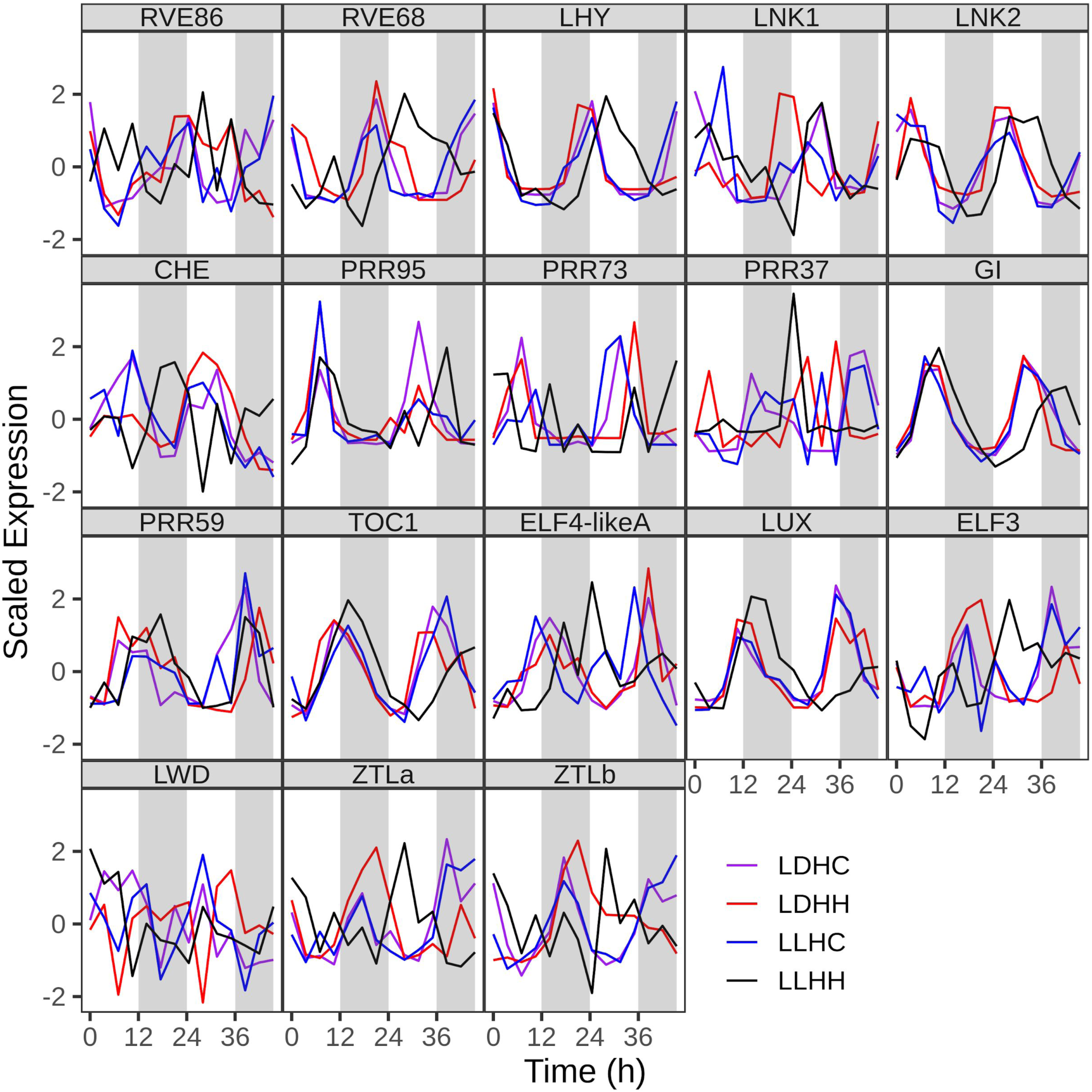
Scaled expression of putative *Brachypodium distachyon* core circadian clock genes in the four time course conditions. Time course names are as described for Figure 1. Source of *B. distachyon* gene identification is described in Supporting Information Table 2.

### Gene expression phase is concentrated to late day and night by external cues and to dawn and dusk by the circadian clock to dawn and dusk

We observed that the majority of all cycling transcripts exhibited an approximately 24 h period. In the presence of any external cue, the vast majority of rhythmic transcripts exhibited a peak expression approximately 8 h after dawn or 8 h after dusk (**Figure 3a**). In contrast to these three conditions, a dramatic shift in expression behavior was observed in the absence of external cues, where the majority of cycling transcripts peaked at dusk with a smaller peak at about 2 h after subjective dawn. This shift in phase could be explained by a lengthening of period observed in constant conditions in contrast to the 24 h period observed under LDHH or LLHC (**Figure 3b**). Transcript expression profiles under LDHC exhibited a double peak shortly before and after 24 h, while in LDHH and LLHC, peak expression occurred after or before dawn, respectively. While constant conditions elicited similar peaks in period distribution, more transcripts exhibited a period of about 27 h suggesting the internal period of Bd21 may be longer than 24 h.

**Figure 3.**
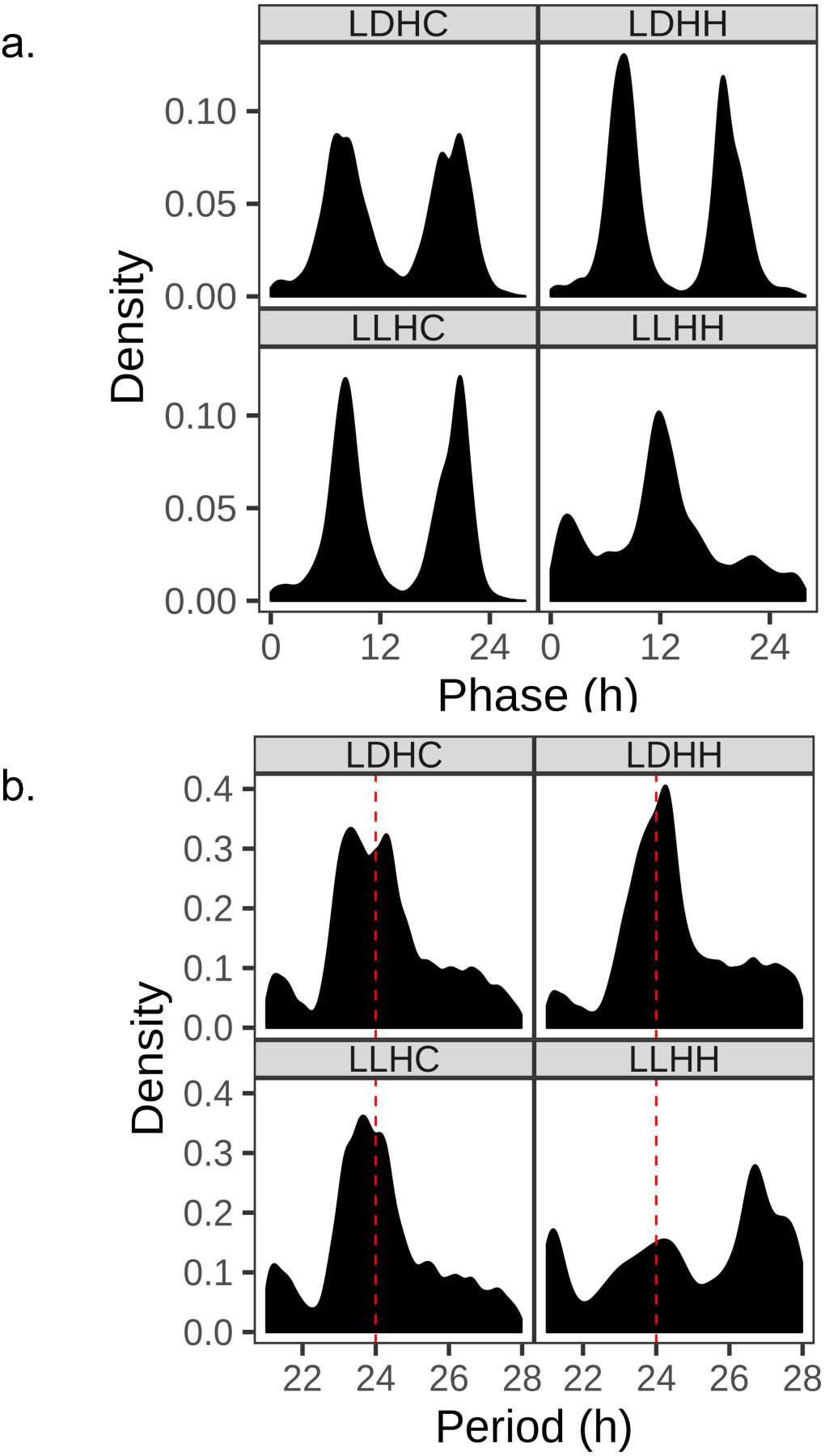
The phase and period of gene expression were primarily conserved in the presence of external cues with a large shift in constant conditions. Distribution of the **(A)** point of maximal expression (phase) and **(B)** period length of cycling transcripts for each condition as determined by MetaCycle. Time course names are as described for Figure 1.

### Thermocycles are the dominant cue driving the 24 h period of circadian clock-regulated genes

To further investigate the relative influence of external cues and the circadian clock on rhythmic gene expression patterns, we analyzed the 306 transcripts found to cycle in all four conditions. As noted before for all rhythmic transcripts, we observed significant effects on period length caused by growth conditions for the 306 cycling genes. Period length was significantly shorter in LLHC and significantly longer in constant conditions (**Table S7**, **Fig. 4a**). While the mean expression values were similar in the presence of external cues, the ranges were not. The interquartile range of the period length distribution was within 1 h of 24 h in LDHC and LLHC and the distributions were significantly different from LDHH and LLHH (**Fig. 4a**). We used hierarchical clustering of period length to group the core cycling transcripts into four non-overlapping sets. We observed that a plurality of transcripts (140) maintained an approximate 24 h period under LDHC, LDHH, and LLHC, but exhibited a significantly longer period under LLHH (**Fig. 4b**, **Fig. S9)**. Forty-seven out of the 306 core cyclic transcripts did not show a significant difference in period between LDHH and LDHC conditions (**Fig. 4d**). Interestingly, 77 transcripts exhibited 24 h periods in LDHC and LLHC, but substantially longer periods in LDHH and LLHH (**Fig. 4e**), while far fewer (42) transcripts exhibited a 24 h period in LDHC and LLHH (**Fig. 4c**). Overall, the patterns show a greater similarity between LDHC and LLHC than LDHC and LDHH. These results suggest that temperature may have a stronger influence than light on transcript periodicity.

**Figure 4.**
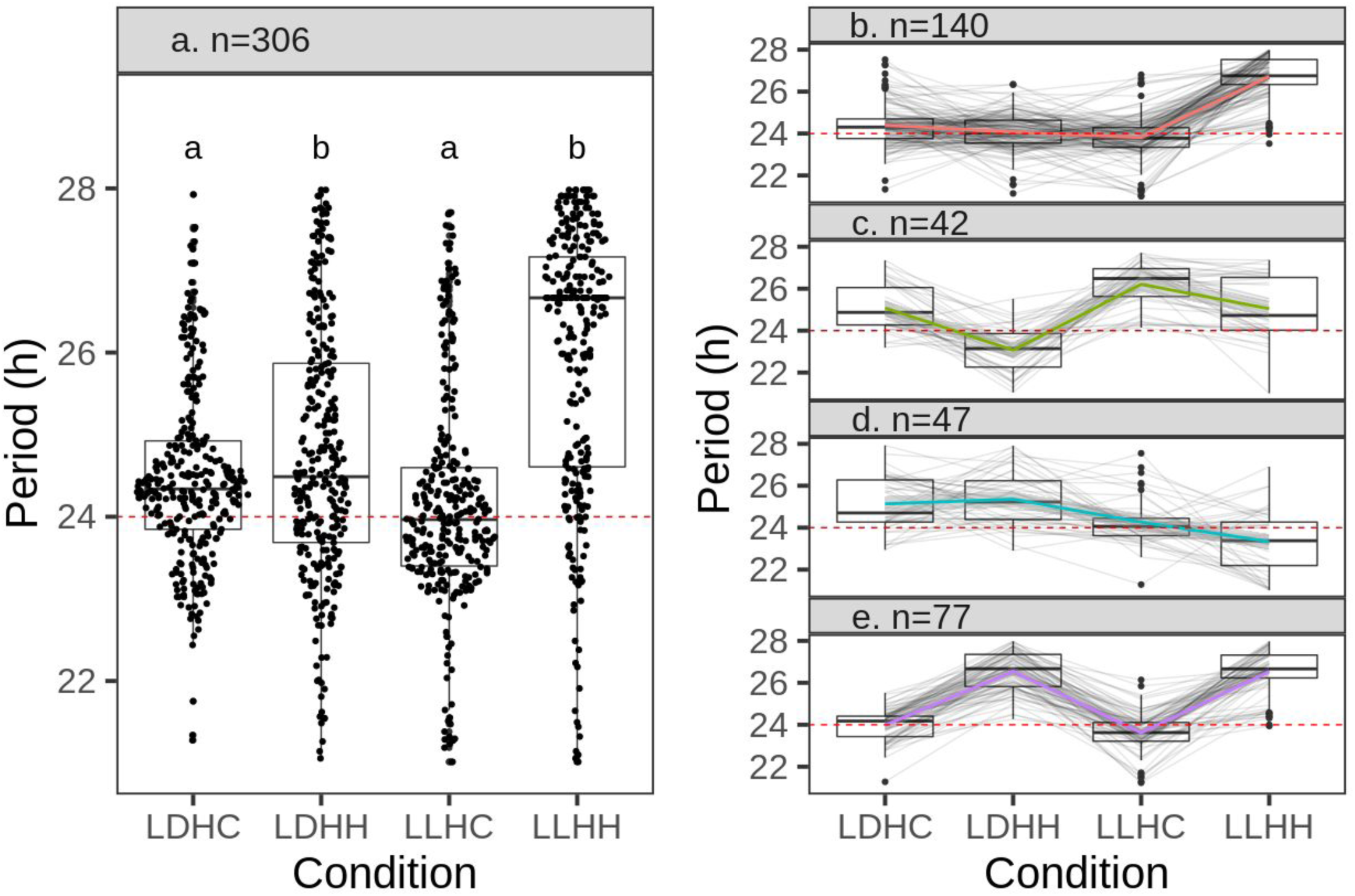
The period length of transcripts rhythmic in all four conditions (a). The period length of the 306 circadian clock-regulated transcripts with a significant period between 21 and 28 h under all four conditions is overlaid on a boxplot to denote the interquartile range of the period data. Each line represents a unique transcript and its period within a condition. Hierarchical clustering was done on normalized period data to group the transcripts into four unique clusters (b-e); n is the number of transcripts in each cluster. The boxes represent the interquartile range, the median value is depicted by a black line, and whisker length is determined as 1.5 times the interquartile range. Letters represent significant (p < 0.01) differences in homoscedasticity as determined by pairwise Fligner-Killeen test of homogeneity of variances. Time course names are as described for Figures 1 and 4.

### Thermocycles are the dominant cue determining amplitude and average expression

Circadian amplitude is the range of expression from the midpoint to the minimum (nadir) and maximum (zenith) for a transcript. Relative amplitude therefore is the ratio between the absolute amplitude and a midline expression for a transcript, allowing for comparison of transcripts with differing expression levels. We observed significant differences in mean relative amplitudes between all four conditions (**Fig. 5a**, **Table S8**). Relative amplitudes of transcripts in LDHC and LLHC conditions were similar (though still statistically distinct, *p* < 0.01 Wilcoxon rank sum test), while cyclic transcripts under LLHH exhibited significantly weaker amplitude (*p* < 0.01). Interestingly, LDHH conditions elicited significantly greater amplitudes among cycling transcripts. Absent external cues, these results suggest that negative regulation of transcript expression is relieved, thus dampening the amplitude and increasing overall expression. We further interrogated the effect of photo- and thermocycles on the amplitude of cycling transcripts by comparing the distribution of expression values. There was a significant increase in mean expression in constant temperature (LDHH and LLHH, **Fig. 5b**). Thus, while LDHH and LLHH conditions have opposing impacts on amplitude, both conditions increased mean expression levels (**Fig. 5c**). As with the period and phase, amplitude and expression levels in LDHC and LLHC conditions were most similar. Taken together, our results indicate that rhythmic gene expression is strongly influenced by thermocycles, with thermocycles alone being sufficient to reproduce the expression patterns observed in the presence of both photo- and thermocycles.

**Figure 5.**
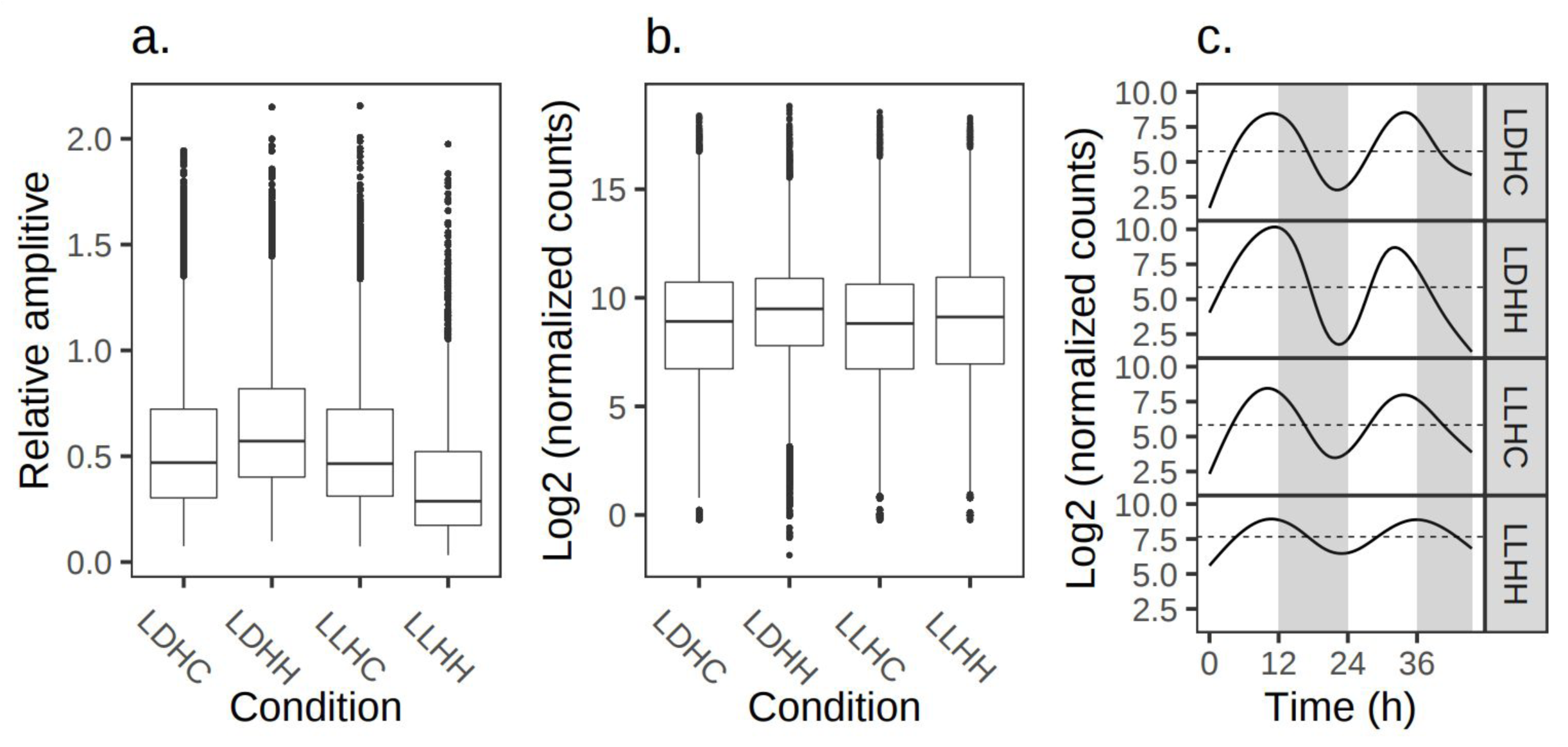
Transcript abundance increased under constant temperatures while photocycles increased relative amplitude. Relative amplitude (**a**) and normalized expression (**b**) of all transcripts rhythmic within each condition. The boxes represent the interquartile range, the median value is depicted by a black line, whisker length is determined as 1.5 times the interquartile range, and outliers are represented by separated dots. (**c**) Representative trendline of all rhythmic transcripts with a phase of approximately 11 h. Is shown to highlight changes in amplitude and expression between conditions. Dotted horizontal line is the mean relative expression level. Box plots and time course names are as described in Figures 1 and 4.

### Gene expression and phase clustering show pathway functions are uniquely or similarly influenced by external cues

To determine whether specific physiological functions are impacted by differences in external cues, we used Pearson correlation coefficients to hierarchically cluster all transcripts that were expressed in all four conditions. Nine clusters were defined based on patterns of expression across each of the conditions (**Fig. S10**). While we applied no filters for diurnal or circadian rhythms, the profiles for each cluster suggest that daily rhythms were important determinants for clustering of transcript expression patterns (**Fig. 6**). The normalized trendlines for LDHC is that of a daily rhythm show that seven of the nine clusters. Those were most often shared in common with LLHC (6) than LDHH (2). Annotations associated with DNA replication and repair or homologous recombination were significantly enriched for genes in cluster 1, which were rhythmic under photocycles and peaked during the day. Terms associated with ethylene signaling, genetic interaction between sugar and hormone signaling, and non-homologous end-joining involved in DNA repair were enriched among transcripts with daytime expression in the presence of thermocycles (clusters 5 and 8). Our results also suggest that genes connected to biosynthesis of secondary metabolites, carbon fixation, and peroxisome activity are expressed during the night and regulated by both photo- and themrocycles to the same time-of-day (cluster 7). Terms associated with mRNA surveillance, which is involved in the quality control and degradation of mRNA, were enriched in cluster 4, which tended to be expressed during the day under thermocycles, but at night under photocycles. The opposite pattern was observed for spliceosome pathway-associated annotations in cluster 9, where expression peaked during the night in the presence of thermocycles and the day under photocycles alone. Genes annotated as aminoacyl-tRNA biosynthesis components were enriched in cluster 6, with peak expression at dusk under all three entrainment conditions. Other clusters of genes that were minimally affected by external cues were enriched for terms associated with processing and transport of RNA, ribosome biogenesis, photosynthetic antenna proteins, and plant-pathogen interactions (clusters 2 and 3). The trendlines in constant conditions did not show a strong circadian waveform in any of the clusters identified. Consistent with our other observations, clusters of genes with rhythmic expression patterns tended to have expression restricted to dawn or dusk, and were more heavily influenced by thermocycles than photocycles (**Figure 6**).

**Figure 6.**
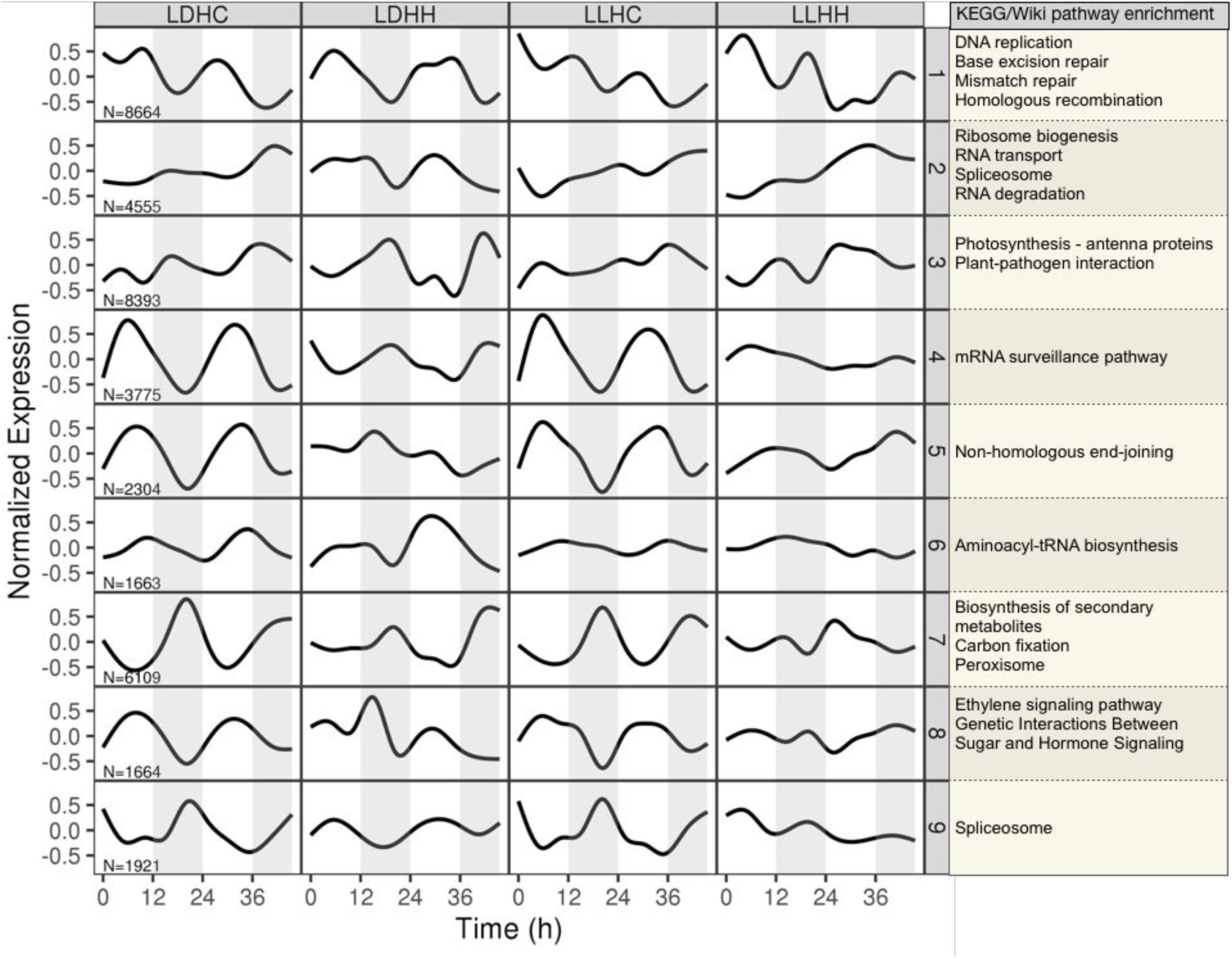
Transcripts associated with specific pathway functions are uniquely or similarly influenced by external cues. Normalized trendline representation of cluster expression by condition along with associated KEGG or Wiki pathway enrichment. Clusters of genes were determined by hierarchical clustering from supporting information fig. 3. Grey rectangles indicate subjective night. N is the number of genes per cluster. Time course names are as described for Figure 1.

Transcripts that were rhythmic in at least one time course were grouped by phase and tested for enrichment of biological process gene ontology (GO) terms (**Fig. S11**). This analysis was meant to identify biological processes that are diurnal or circadian clock controlled, and whether they exhibit variable expression depending on diurnal condition. Notably, secondary cell wall biosynthesis (including terms associated with ‘secondary cell wall biogenesis’, and ‘lignin biosynthetic process’) appeared to be thermocycle regulated from this analysis (**Fig. S11**, **Fig. 7**). This observation is also consistent with gene expression profiles exhibited by hierarchical cluster 7 (**Figure 6**) and suggests strong coordinated action of thermocycles in regulating control of plant structural properties to the night. GO terms associated with pathogen defense pathways appeared in both thermocycle (response to fungus, jasmonic acid biosynthetic process, defense response to bacterium) and photocycle (response to chitin, response to jasmonic acid stimulus, defense response) driven conditions independently, but the same terms were not associated with all conditions. This is consistent with our hierarchical clustering observations that found pathogen responses were not strongly driven by any one condition (cluster 3; **Fig. S11**). Similar to our observations from hierarchical clustering of gene expression profiles, GO term enrichment analysis suggests that photocycles elicited rhythmic expression of gene sets related to cellular processes, such as DNA replication, translational elongation, and tricarboxylic acid cycle. These groups tended to be expressed during the day (**Fig. S11**). The only terms to appear across all conditions are ‘response to heat’, ‘photosynthesis light harvesting’, and ‘photosynthesis’.

**Figure 7.**
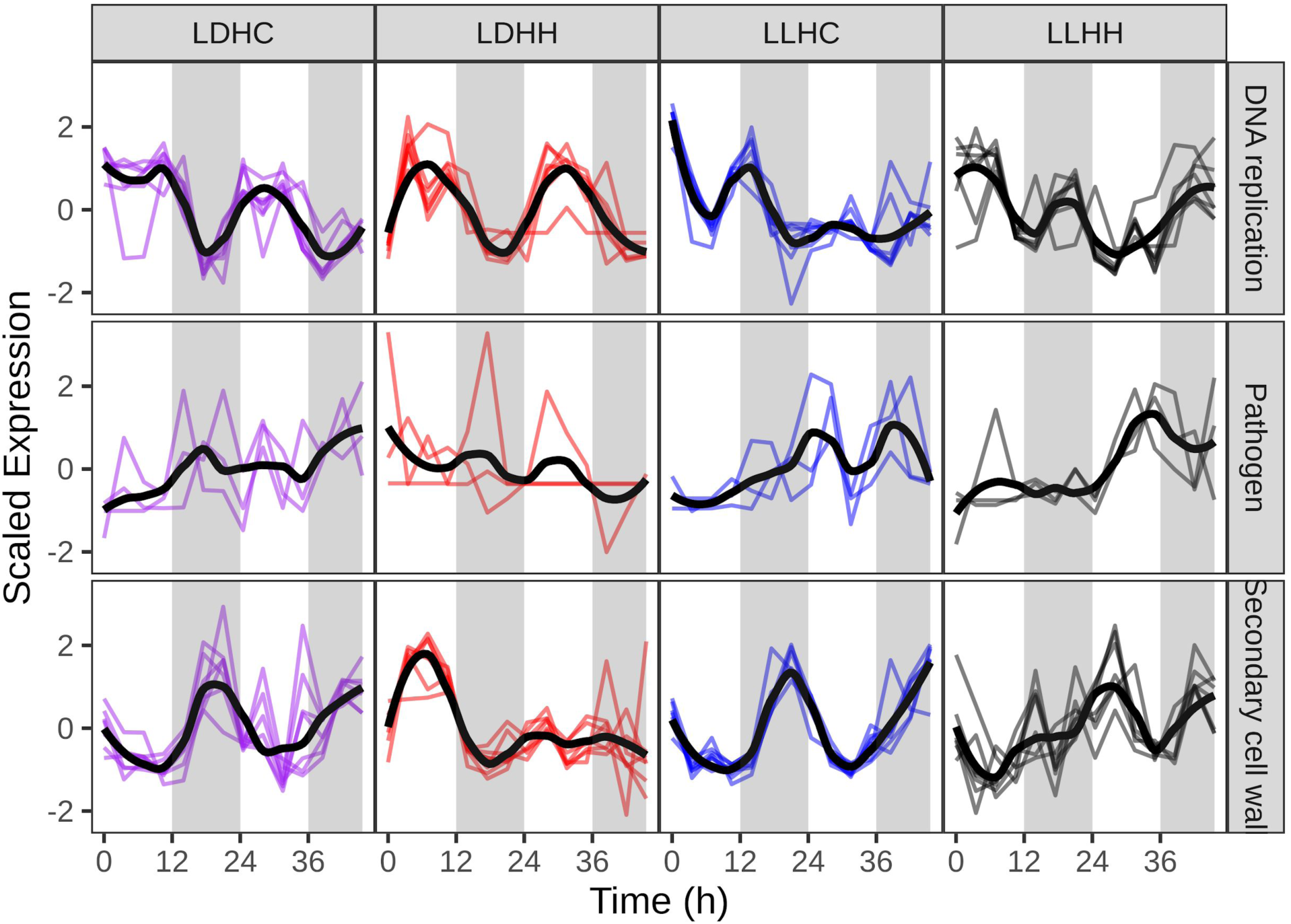
Scaled expression of representative transcripts associated with the given gene ontology term. Black line is the trendline for the transcripts shown. Colored lines are individual transcripts. Gray vertical bars represent subjective night. Time course names are as described for Figure 1.

### Time-of-day specific *cis*-regulatory sequences

It has been shown that some *cis*-regulatory sequences controling time-of-day expression are conserved across monocots and dicots (Michael *et al*., 2008b; Filichkin *et al*., 2011). Three *cis*-element modules made up of the morning element (ME:CCACAC), evening element (EE:AAATATCT), and the telobox (TBX:AAACCCT) along with their associated elements, including CCA1-binding site (CBS:AAAATCT), G-box (CACGTG), GATA, starch box (SBX:AAGCCC) and protein box (PBX:ATGGGCCG) (Michael *et al*., 2008b,a; Filichkin *et al*., 2011). To identify known and novel time-of-day-specific *cis*-elements, we used the ELEMENT program to perform a search for 3-8 bp sequences. We identified 113, 86, 327 and 11 significantly overrepresented *cis*-elements under LDHC, LLHC, LDHH and LLHH, respectively, including ME, EE, G-box core (ACGT), TBX, PBX, and SBX (**Fig. 8**) (Michael *et al*., 2008b). However, TBX, SBX, and PBX elements were only significantly overrepresented in LDHH conditions. Inspection of all significantly overrepresented 3-8 bp sequences revealed a clear pattern that conditions with thermocycles are alike and distinct from those with photocycles alone (**Fig. S12**). These results suggest that thermocycles override the impact of photocycles on gene expression regulation, consistent with our previous results that thermocycles play a distinct and dominant role in coordinating time-of-day expression.

**Figure 8.**
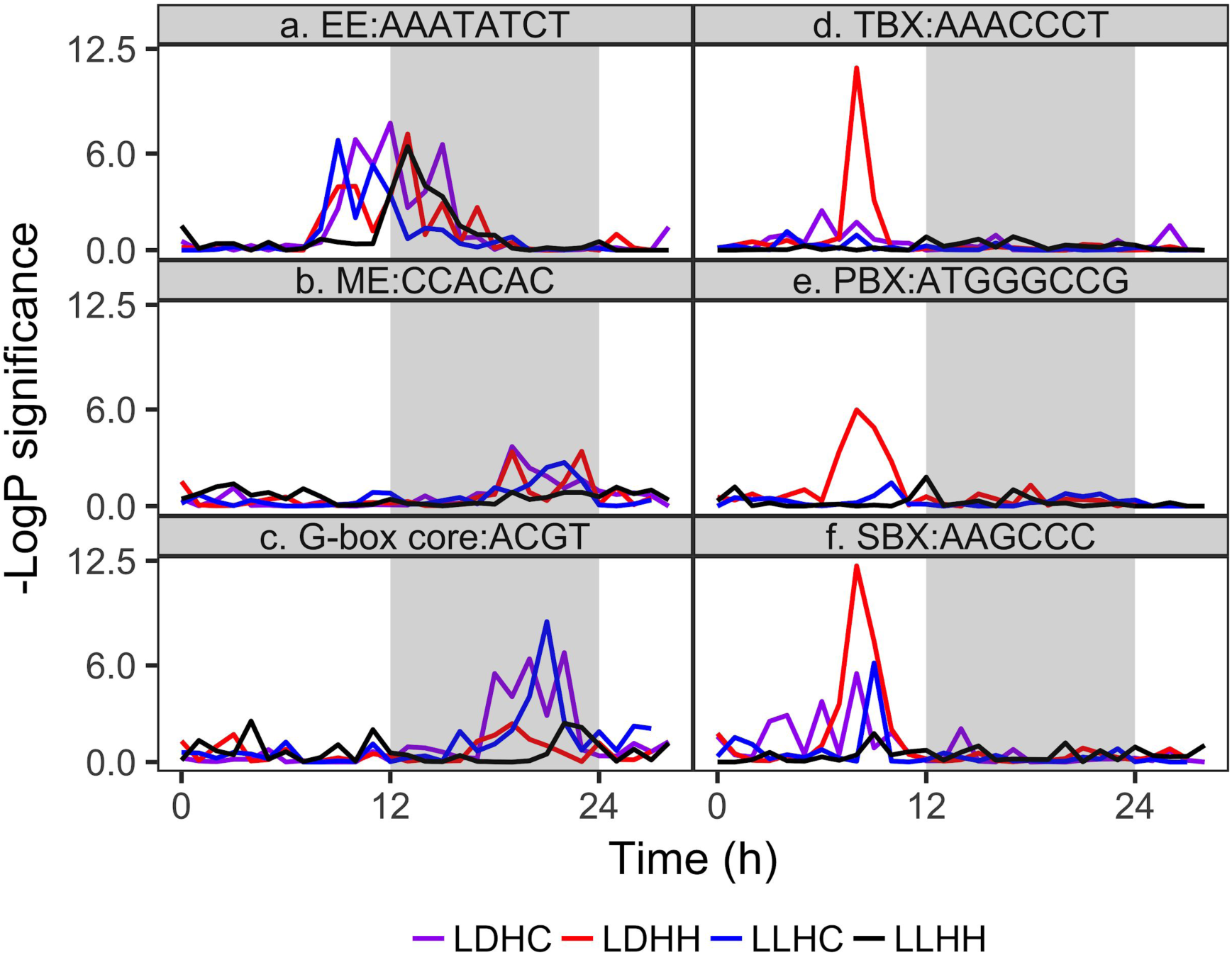
Time-of-day specific diurnal and circadian *cis*-elements. Negative log P values for known diurnal and circadian cis-elements (a) evening element (EE), (b) morning element (ME), (c) G-box core, (d) telobox (TBX), (e) protein box (PBX) and (f) starch box (SBX) were plotted across the day to identify the time of day they were overrepresented in *cis*-regulatory region of genes that cycle with that specific phase.Time course names are as described for Figure 1.

DNA affinity purification sequencing (DAP-seq) is a powerful method to identify transcription factor binding sites *in vitro* (O’Malley *et al*., 2016). We leveraged DAP-seq binding site motifs from *A. thaliana* to link time-of-day motifs to their putative transcription factors in the *B. distachyon* time courses. We identified 291 (out of 477 total DAP-seq motifs) significantly enriched motifs, including the core diurnal elements EE, GATA, TBX and G-box (**Fig. 9**, **Fig. S13**). In contrast to ELEMENT, this approach enabled us to identify larger, split elements like VNS (VND/NST/SND, CTTNNNNNNAAG, (Olins *et al*., 2018)), which was enriched in constant conditions and thermocycles, suggesting that it is very likely part of circadian clock regulation. In addition, we observed enrichment for the CACT motif in the morning in LDHC and LDHH. This element is associated with C2H2 family transcription factors (O’Malley *et al*., 2016), and is thus a potentially photocycle responsive motif (**Fig. S14**). The CRT/DRE motif (CCGAC, (Stockinger *et al*., 1997; Liu *et al*., 1998)), which is bound by CBF type AP2-EREBP proteins, is enriched in midday- and nighttime-expressed transcripts. In addition to the CRT/DRE motif, three other motifs were enriched in the nighttime: C2C2-Dof binding AAAAAGG (Ko *et al*., 2016), WRKY-binding W-box GTCAA (Ulker & Somssich, 2004), and a bZIP binding HEX element TGACGT (Schindler *et al*., 1992). The C2C2-Dof site and the W-box were enriched in all diurnal conditions but not constant conditions. The bZIP site and CRT/DRE motif were enriched in constant conditions and in the presence of thermocycles.

**Figure 9.**
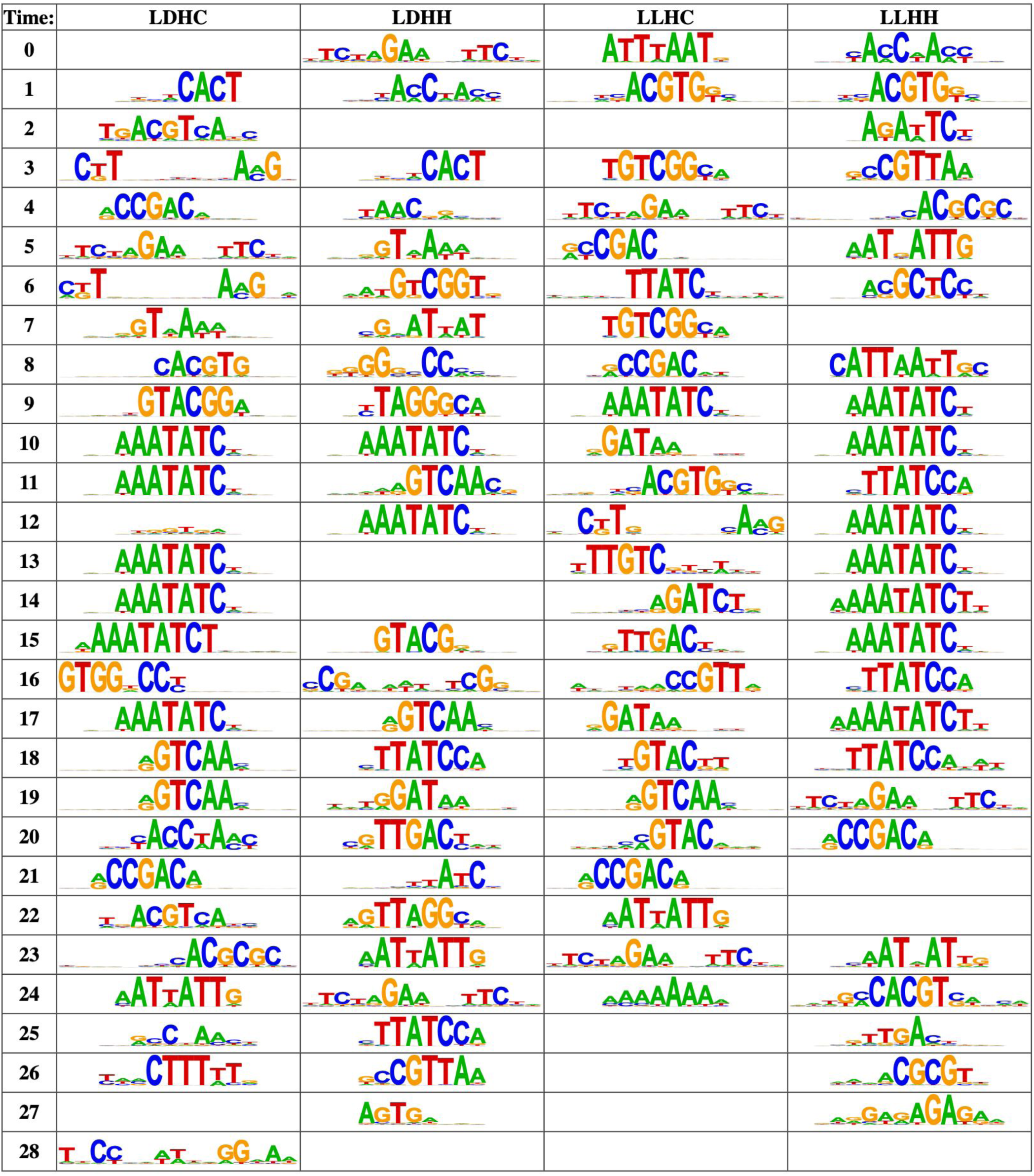
*Cis*-regulatory sequences most overrepresented at each hour in each time course. Each nucleotide sequence is a position probability matrix motif derived from DNA-affinity purification sequencing and identified as enriched in diurnal (LDHC, LDHH, and LLHC) or constant (LLHH) conditions. The height of the letter at each position is proportional to the probability of a given nucleotide. Time course names are as described for Figure 1.

## DISCUSSION

Daily rhythms in transcript abundance are either a direct response to predictable environmental cues including photocycles, thermocycles, and humidity cycles, or they are generated by the circadian clock. Most estimates of daily gene expression patterns have been determined for protein-coding genes using gene or genome tiling microarrays (Edwards *et al*., 2006; Covington *et al*., 2008; Michael *et al*., 2008b; Hazen *et al*., 2009; Khan *et al*., 2010; Filichkin *et al*., 2011). More recently, RNA-seq was applied to measure diurnal or circadian-associated transcript expression in sugarcane, *S. viridis*, *Sedum album*, lettuce, and Douglas fir (Hotta *et al*., 2013; Higashi *et al*., 2016; Cronn *et al*., 2017; Huang *et al*., 2017; Wai *et al*., 2019). Estimates for the proportion of protein-coding genes that are controlled by the circadian clock generally range between 6 and 15%. Species with values well outside this range include *S. viridis* with 1.2% when entrained in thermocycles alone and 33% in sugarcane (Hotta *et al*., 2013; Huang *et al*., 2017). Here we report that 3.6% of all *B. distachyon* transcripts are regulated by the circadian clock, which is small relative to other species. It is worth noting that different analyses applied to the same dataset can identify different rhythmic gene sets (Khan *et al*., 2010; Hotta *et al*., 2013; Hughes *et al*., 2017), potentially due to differences in entrainment conditions, as well as other environmental parameters including light intensity and temperature range. Thus, comparisons of the proportion of circadian-clock-regulated genes among species should be made cautiously, with consideration of the method used to measure and analyze transcript abundance and entrainment conditions. Keeping these limitations in mind, we observed a comparable number of rhythmic genes in the presence of external cues and a relatively low number in constant conditions between *B. distachyon* and other species. This observation suggests that the circadian clock has a relatively minor role in *B. distachyon* gene regulation.

In the presence of external cues, peak gene expression was concentrated at the end of the day and night. In constant conditions there was a dramatically different pattern, with dusk as the most represented phase, followed by subjective dawn. A concentration of genes that peak in expression at the end of the day or at night in some combination of photo- and thermocycles was similarly observed in *A. thaliana*, poplar, rice, and Douglas fir (Covington *et al*., 2008; Michael *et al*., 2008b; Filichkin *et al*., 2011; Cronn *et al*., 2017)(Michael *et al*., 2008b; Hazen *et al*., 2009). In maize and sugarcane, the most abundant phases were centered around dawn and dusk (Khan *et al*., 2010; Hotta *et al*., 2013), similar to the phase expression profile in this study. However, only constant conditions were reported for maize (Khan *et al*., 2010) and sugarcane (Hotta *et al*., 2013) and not similarly described for rice (Filichkin *et al*., 2011). In barley and rice, the expression of putative clock genes was measured under entrainment and constant conditions with no change in period or phase noted between conditions (Murakami *et al*., 2007; Filichkin *et al*., 2011; Campoli *et al*., 2012). Thus, a similar phase shift was neither tested nor observed in several grass species.

The change in phase we observed may be a result of the striking period difference observed in constant conditions. Our analysis was restricted to genes with periods ranging from 21 to 28 h. A vast majority of rhythmic transcripts cycled with a period between 23 and 25 h. While some genes cycled with a similar period in constant conditions, the most abundant period for genes in constant conditions was much longer, with a range between 26 to 28 h. A meta-analysis of 11 different *A. thaliana* time courses did not report a lengthening of period in constant conditions for the Col-0 accession (Michael *et al*., 2008b). Similarly, *S. viridis* genes that were rhythmic under constant conditions exhibited an approximate 24 h period. In contrast, variation in period length has been measured in a number of other species, and in many cases mutations conferring clock-related phenotypes in crop plants have been identified (Bendix *et al*., 2015; Greenham *et al*., 2017)(Michael *et al*., 2003; Dodd *et al*., 2005; Srivastava *et al*., 2019). Also, light intensity is known to influence period length, and has been shown to result in changes at very low fluences (Somers *et al*., 1998; Oakenfull & Davis, 2017). Thus, we cannot rule out an effect of genetic variation or light intensity in our data set without further testing. The 27 h period length estimation in constant conditions in *B. distachyon* may reflect the specific period of the accession tested, but within the range of natural genetic variation observed in several species.

A number of factors are known to alter amplitudes of gene expression. Cold dampens amplitude in *A. thaliana* and a local infection of the bacterial pathogen *Pseudomonas syringae* as well as salicylic acid or H_2_O_2_ results in a systematic decrease in amplitude (Bieniawska *et al*., 2008; Li *et al*., 2018). Several *A. thaliana* circadian clock mutants exhibit a decrease in amplitude up to a total loss of circadian function (Nagel & Kay, 2013). Often, these changes in amplitude cause an increase in average gene expression because the nadir is not as low. Our results suggest that a self sustaining circadian clock provides only a small portion of the daily repression that results in robust amplitude and waveform.

Circadian clock genes and their expression behavior are largely conserved between grasses and eudicots (McClung, 2010; Campoli *et al*., 2012Mockler *et al*., 2007; Michael *et al*., 2008b; Filichkin *et al*., 2011; Wai *et al*., 2019). Several genes, namely *BdRVE86*, *BdLNK1*, *BdCHE*, *BdPRR73*, *BdELF4-likeA*, and *BdELF3* were not detected as significantly circadian clock regulated in this study. Under entrainment conditions, the average clock gene cycled with a nearly 24 h period, but this period lengthened to greater than 26 h in constant conditions. It may be that the absence of rhythmic expression of one or more circadian clock genes resulted in a substantially slower circadian clock in *B. distachyon*. Mutations in all of the arrhythmic genes can result in a long period or arrhythmic expression in *A. thaliana* (Nagel & Kay, 2013). ELF3 is a particularly interesting candidate as naturally occurring allelic variants in *A. thaliana* are linked to poor rhythms (Box *et al*., 2015).

Analysis of the *cis*-regulatory regions of rhythmic genes with similar time-of-day expression identified sequence motifs that are candidates for functional determinants of that expression behavior. In spite of the relatively small number of circadian-clock-regulated transcripts identified in our study, we recovered a robust signature for the evening element under all four conditions. The G-box, a morning-enriched motif in other systems (Zdepski *et al*., 2008; Filichkin *et al*., 2011), was enriched in both morning and evening expressed genes in thermocycles and constant conditions. We also observed enrichment of the CBF binding site among evening-phased genes in all conditions except LDHH. The CBF family of transcription factors is strongly associated with the circadian clock, providing temperature input by directly binding the *LUX* promoter (Chow *et al*., 2014; Gierczik *et al*., 2017). Neither the G-box nor the CRT element was overrepresented under photocycles alone. We further identified several motifs not previously described as having a role in diurnal gene expression or circadian clock output. These include the VNS, W-box, and Dof binding motif. The VNS motif featured heavily in secondary cell wall development, and is a consensus sequence bound by the VND, NST, and SND NAC transcription factors (Olins *et al*., 2018). Consistent with our observation of rhythmic secondary cell wall genes, this motif was enriched in both diurnal and constant conditions. The W-box is a binding target for some WRKY transcription factors, which have been shown to interact with the circadian clock to help regulate senescence (Kim *et al*., 2018). ChIP-seq experiments have revealed the Dof-binding motif, AAAAAGG, to be a target of ZmCCA1 and LHY (Ko *et al*., 2016; Adams *et al*., 2018). Similarly, several CDF transcription factors are clock regulated and play a crucial role in photoperiodic timing (Goralogia *et al*., 2017). The HEX motif is associated with bZIP-binding in plants and represent a possible source of entrainment for the circadian clock through metabolic adjustments in response to sugar availability (Schindler *et al*., 1992; Frank *et al*., 2018). Overall, we identified a number of sequence motifs not previously associated with rhythmic gene expression, among them possible drivers of thermocycle rhythms.

Rhythmic growth in plants has been observed across numerous species, and depends on both internal and external cues (Walter *et al*., 2009; Farre, 2012). In *A. thaliana*, the rate of hypocotyl elongation is greatest at the end of the night and is heavily influenced by photoreceptors and the circadian clock (Dowson-Day & Millar, 1999; Nozue *et al*., 2007). In grasses, temperature is the dominant cue that determines the relative rate of cell division and elongation (Watts, 1971; Poiré *et al*., 2010). Previously, we reported that the rate of *B. distachyon* leaf elongation was regulated by thermocycles (Matos *et al*., 2014). Here we observed that genes associated with secondary cell wall biosynthesis, including *CELLULOSE SYNTHASE A4*/*7*/*8* and several lignin pathway genes (Coomey & Hazen, 2016), were regulated by thermocycles with peak expression occurring at night. Rhythmic coexpression of lignin and cell-wall-related genes has been observed in maize and *A. thaliana*, but often with different timing or following different cues (Michael *et al*., 2008a; Khan *et al*., 2010). Indeed, in *A. thaliana*, carbon status and light influence lignin gene expression and hypocotyl lignification (Rogers *et al*., 2005). It is interesting to note that our gene expression results clearly show that expression of secondary-wall-related genes follows thermocycles, but is antiphasic to reported maximal elongation rates (Matos *et al*., 2014). This suggests that rhythmic elongation under warm temperatures may be followed by rhythmic wall thickening under nighttime temperatures.

In summary, both internal and external cues caused pervasive circadian rhythms in *B. distachyon*. The total effect on rhythmic gene expression was similar between photo- and thermocycles, and more than 21% of genes had rhythmic expression in the presence of both cues. Thermocycling appears to be the prevailing signal. The overlap was greatest between genes that commonly cycled in LDHC and LLHC. Rhythmic gene expression under thermocycles had a more narrow distribution and a period closer to 24 h for genes rhythmic in all four conditions. Overall expression level and relative amplitude was most similar between LDHC and LLHC. Hierarchical clustering revealed pathway enriched groups of genes more commonly expressed between LDHC and LLHC and clustering of gene sets based on phase of expression revealed distinctly synchronized pathways between thermocycles and photocycles. The relative importance of thermocycles is consistent with the observation that grasses are uniquely void of elongation growth rhythms in constant conditions or in the presence of photocycles alone (Poiré *et al*., 2010; Matos *et al*., 2014).

## AUTHOR CONTRIBUTIONS

BC, SH, and SK conceived and designed the study. BC, MR, and TD acquired the data. KM, BC, CY, JC, NH, TM, and SH analyzed and interpreted the data. KM, BC, JC, TM, and SH drafted the manuscript.

## SUPPORTING INFORMATION

Fig. S1. Distribution of read counts per time point between conditions.

Fig S2. Comparison of period and phase estimates for all cycling transcripts with a phase estimate of approximately 12 h.

Fig. S3. Distribution of Metacycle p-value assignments across conditions.

Fig. S4. Relative expression of suggested controls for single-gene PCR quantitation. Fig. S5. Phylogenetic analysis of *A. thaliana RVE* genes *AtRVE6* and *AtRVE8*.

Fig. S6. Phylogenetic analysis of *A. thaliana LWD* genes, *AtLWD1* and *AtLWD2*. Fig. S7. Phylogenetic analysis of *A. thaliana LNK* genes *AtLNK1* and *AtLNK2*.

Fig. S8. Comparison of *A. thaliana* and *B. distachyon* photoreceptor expression profiles.

Fig. S9. 95% family-wise confidence level of mean cluster periods from post-hoc Tukey’s honest significant differences test.

Fig. S10. Heatmap showing z-score normalized expression of all transcripts expressed in all four conditions, segregated into hierarchical clusters.

Fig. S11. Gene ontology terms overrepresented in each phase across the four conditions.

Fig. S12. Heatmap of significant 3-8 bp elements across conditions.

Fig. S13. Complete list of cis-regulatory sequences overrepresented for each phase in each of the conditions.

Fig. S14. Non-redundant set of enriched DAP-seq motifs and highest likelihood *A. thaliana* transcription factor matches.

Table S1. Total number of trimmed reads and percent mapped for each sequenced library.

Table S2. Protein sequences used for phylogenetic analysis.

Table S3. List of previously recommended control genes for single-gene PCR quantitation and their associated coefficient of variance.

Table S4. List of nine *B. distachyon* transcripts with minimal expression change between conditions or time points.

Table S5. List of *B. distachyon* circadian clock genes and their *A. thaliana* ortholog, along with associated period and phase.

Table S6. Putative *B. distachyon* photoreceptor gene primary transcripts and aliases identified previously and by amino acid sequence comparison with *A. thaliana*.

Table S7. Summary of pairwise Wilcoxon rank sum significance tests performed for period length comparisons.

Table S8. Summary of pairwise Wilcoxon rank sum significance tests performed for relative amplitude comparisons.

## Supplemental material

**Supporting Information Table 1.** Total number of trimmed sequence reads per time point and the percentage that mapped to Bd21 v3.1 transcriptome. RNA-seq was performed on all samples with an average read length of 70 bp. Raw reads were first processed with TrimGalore and FastX Trimmer to remove low quality reads. LDHC, photo- and thermocycles; LDHH, photocycles and constant temperature; LLHC, thermocycles and constant light; LLHH, constant light and warm temperature.

| Time (h) | LDHC |  | LDHH |  | LLHC |  | LLHH |  |
| --- | --- | --- | --- | --- | --- | --- | --- | --- |
|  | Reads | Percent Mapped | Reads | Percent Mapped | Reads | Percent Mapped | Reads | Percent Mapped |
| 0.0 | 34,227,208 | 94.48 | 35,742,006 | 97.75 | 40,051,669 | 97.96 | 41,236,428 | 96.78 |
| 3.5 | 35,154,998 | 94.78 | 48,487,323 | 97.86 | 37,518,944 | 98.7 | 39,289,113 | 97.5 |
| 7.0 | 33,914,214 | 94.63 | 44,882,338 | 97.82 | 38,623,206 | 98.56 | 33,929,870 | 97.32 |
| 10.5 | 34,489,682 | 94.83 | 48,455,805 | 98.16 | 39,693,845 | 98.5 | 39,963,994 | 97.93 |
| 14.0 | 36,778,363 | 94.59 | 41,426,536 | 98.36 | 39,244,249 | 98.51 | 37,832,873 | 97.63 |
| 17.5 | 33,906,037 | 94.11 | 110,448,634 | 97.97 | 36,885,205 | 98.77 | 36,486,074 | 98.02 |
| 21.0 | 41,607,392 | 95.15 | 18,925,532 | 94.74 | 37,676,926 | 96.11 | 38,278,956 | 98.16 |
| 24.5 | 39,515,647 | 98.59 | 17,350,554 | 95.9 | 38,352,753 | 98.22 | 37,491,784 | 97.99 |
| 28.0 | 40,636,672 | 98.58 | 16,526,350 | 95.06 | 33,838,170 | 97.99 | 38,780,829 | 98.56 |
| 31.5 | 40,356,148 | 98.41 | 18,991,591 | 96.65 | 35,994,479 | 98.34 | 40,582,785 | 98.61 |
| 35.0 | 38,675,103 | 98.66 | 21,406,132 | 94.62 | 34,695,919 | 98.59 | 38,521,775 | 98.72 |
| 38.5 | 39,094,086 | 98.73 | 22,334,225 | 92.45 | 38,871,544 | 98.28 | 41,319,197 | 98.93 |
| 42.0 | 33,613,347 | 98.43 | 28,354,992 | 95.23 | 37,907,439 | 97.12 | 39,499,304 | 98.82 |
| 45.5 | 36,555,043 | 98.65 | 10,626,906 | 95.75 | 36,998,110 | 97.14 | 38,625,672 | 98.64 |

**Supporting Information Table 2.**
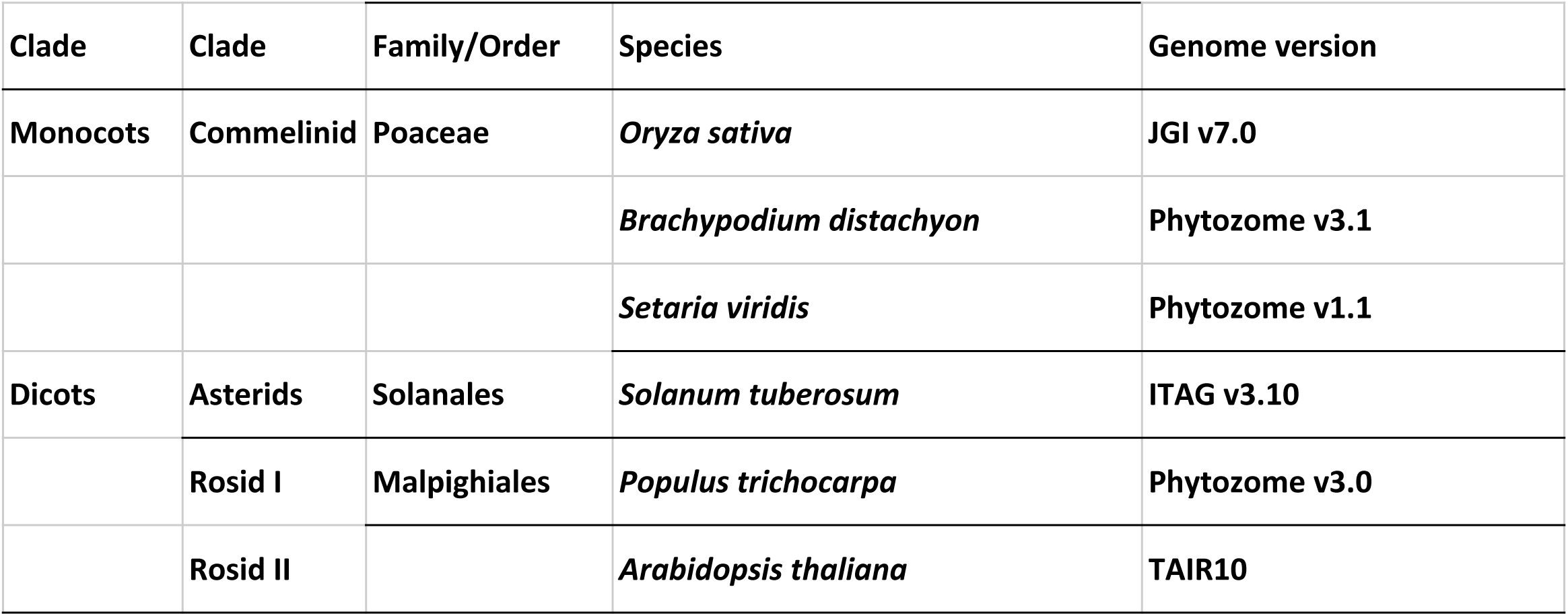
Phylogeny proteins, separate file.

| Clade | Clade | Family/Order | Species | Genome version |
| --- | --- | --- | --- | --- |
| Monocots | Commelinid | Poaceae | *Oryza sativa* | JGI v7.0 |
|  |  |  | *Brachypodium distachyon* | Phytozome v3.1 |
|  |  |  | *Setaria viridis* | Phytozome v1.1 |
| Dicots | Asterids | Solanales | *Solanum tuberosum* | ITAG v3.10 |
|  | Rosid I | Malpighiales | *Populus trichocarpa* | Phytozome v3.0 |
|  | Rosid II |  | *Arabidopsis thaliana* | TAIR10 |

**Supplemental Table 3.**
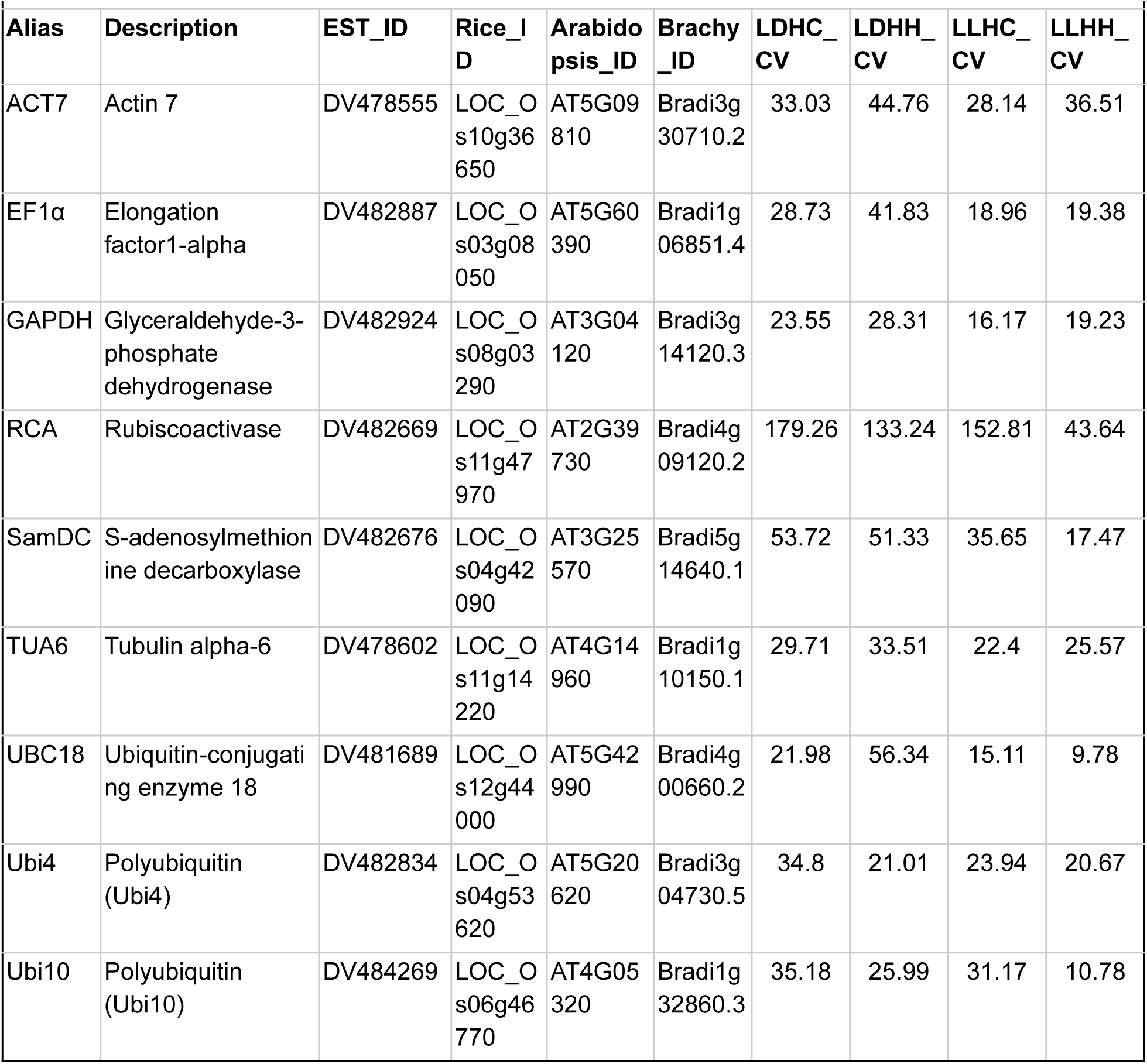
Table of housekeeping genes as recommended by (Hong et al. 2008). Coefficient of variance for all genes is shown as CV in each condition. Time course names are as described in Figure 1.

**Supporting Information Table 4.** List of nine *B. distachyon* transcripts with minimal expression change between conditions or time points. Each listed transcript is in the top two percentile for coefficient of variance for each of the four conditions.

| Alias | Description | Rice_ID | Arabidopsis_ID | Brachy_ID | LDHC_CV | LDHH_CV | LLHC_CV | LLHH_CV |
| --- | --- | --- | --- | --- | --- | --- | --- | --- |
| ATRAB7B | RAB GTPase homolog G3F | LOC_Os05g44050.1 | AT3G18820.1 | Bradi2g19756.1 | 10.75348903 | 11.15410056 | 9.271679696 | 8.087304627 |
|  |  | LOC_Os02g10050.1 | AT2G22370.1 | Bradi3g06860.1 | 11.50338629 | 9.618858702 | 10.43579708 | 8.071010146 |
|  | Endosomal targeting BRO1-like domain-containing protein | LOC_Os10g35250.1 | AT1G15130.1 | Bradi3g29880.1 | 10.89108066 | 7.31915461 | 10.26570475 | 5.964732371 |
|  | Serine carboxypeptidase S28 family protein | LOC_Os10g36760.1 | AT4G36195.1 | Bradi3g30640.1 | 8.085289622 | 11.09028911 | 9.079985663 | 5.907313041 |
|  |  | LOC_Os11g03380.1 | AT3G13410.1 | Bradi4g43990.1 | 8.874407152 | 10.00288149 | 8.512653509 | 10.12325971 |
|  | Peptidase S24/S26A/S26B/S26C family protein | LOC_Os04g01240.1 | AT2G31140.1 | Bradi5g02390.1 | 8.872811828 | 11.13538202 | 8.314756559 | 8.243739656 |
|  | Glutathione S-transferase family protein | LOC_Os04g17050.1 | AT5G42150.1 | Bradi5g04550.1 | 6.809561668 | 10.9006132 | 9.171676531 | 8.289582374 |
| ATPIS1 | phosphatidylinositol synthase 1 | LOC_Os02g03110.2 | AT1G68000.1 | Bradi5g22340.1 | 9.812465271 | 11.21326709 | 11.10497261 | 7.126786259 |
|  |  | LOC_Os04g58560.1 | AT2G44140.1 | Bradi5g26480.1 | 7.651585894 | 9.795829623 | 11.02811865 | 6.01837538 |

**Supporting Information Table 5.** Putative *Brachypodium distachyon* circadian clock gene primary transcripts and aliases identified previously and by amino acid sequence comparison with *Arabidopsis thaliana* and sorted by average phase in the presence of external cues.

| Gene | *B. distachyon* ID | *A. thaliana* ortholog | Time of activity in *A. thaliana* | Period (h) |  |  |  | Phase (h) |  |  |  |
| --- | --- | --- | --- | --- | --- | --- | --- | --- | --- | --- | --- |
|  |  |  |  | LDHC | LDHH | LLHC | LLHH | LDHC | LDHH | LLHC | LLHH |
| *RVE86* | Bradi1g32330.1<sup>a</sup> | At3g09600 | Midday to afternoon | 23.6* | - | 27.1 | - | 21.2 | - | 20.5 | - |
| *RVE68* | Bradi3g51757.6<sup>a</sup> | At5g52660 | Midday to afternoon | 23.7 | 25.9 | 21.5 | 26.4 | 2.8 | 23.4 | 4.7 | 2.8 |
| *CCA1* | Bradi3g16515.1<sup>b</sup> | At1g01060 | Dawn | 24.5 | 25.7 | 24.1 | 26.7 | 23.3 | 23.1 | 22.6 | 2.3 |
| *LNK1* | Bradi1g60846.1<sup>a</sup> | At5g64170 | Midday to afternoon | 25.9 | 24.4* | 23 | - | 2.7 | 4.8 | 4.5 | - |
| *LNK2* | Bradi2g13610.2<sup>a</sup> | At3g54500 | Midday to afternoon | 24 | 23.9 | 23.8 | 26.3 | 2.6 | 4.5 | 2.6 | 6.5 |
| *CHE* | Bradi3g60352.1<sup>b</sup> | At5g08330 | Evening | 23.1 | 24.2 | - | - | 7.8 | 5.9 | - | - |
| *PRR95* | Bradi4g36077.1<sup>b</sup> | At2g46790 | Morning/afternoon | 23.8 | 23.5* | - | 26.4 | 8.7 | 7.1 | - | 11.2 |
| *PRR73* | Bradi1g65910.1<sup>b</sup> | At5g02810 | Midday/evening | 26.6 | 21.5* | - | - | 8.4 | 7.5 | - | - |
| *PRR37* | Bradi1g16490.6<sup>b</sup> | At5g60100 | Midday/evening | 24.5 | 21 | 25.3* | 24.1 | 9.7 | 10.6 | 8.1 | 12.2 |
| *GI* | Bradi2g05226.1<sup>b</sup> | At1g22770 | Evening | 24.4 | 23.3 | 24.9 | 27.6 | 10.1 | 9.6 | 9.7 | 11.6 |
| *PRR59* | Bradi4g24967.1<sup>b</sup> | At5g24470 | Morning/afternoon | 26.5 | - | - | 25.4 | 10.5 | - | - | 16.1 |
| *TOC1* | Bradi3g48880.1<sup>b</sup> | At5g61380 | Evening | 23.7 | 24.2 | 23.1 | 26.7 | 13 | 11.7 | 14.1 | 16.5 |
| *ELF4-li keA* | Bradi4g29580.1<sup>c</sup> | At2g40080 | Evening | 24 | 24.5* | 23.3 | - | 14.9 | 15 | 10.9 | - |
| *LUX* | Bradi2g62067.1<sup>d</sup> | At3g46640 | Evening | 23.5 | 24.8 | 23.3 | 27.8 | 13.9 | 13.6 | 13.4 | 17.9 |
| *ELF3* | Bradi2g14290.1<sup>b</sup> | At2g25930 | Evening | 22.8 | 26.9 | - | - | 17.7 | 19.5 | - | - |
| *LWD* | Bradi3g44580.1<sup>a</sup> | At1g12910 | Morning | - | - | - | - | - | - | - | - |
| *ZTLa* | Bradi1g33610.2<sup>a</sup> | At5g57360 | Night | 22.8* | 25.4 | 24.4* | 26.7* | 18.5 | 19.2 | 16.9 | 2 |
| *ZTLb* | Bradi3g04040.1<sup>a</sup> | At5g57360 | Night | 21.5 | - | 25.7 | - | 18.6 | - | 17.9 | - |

**Supporting Information Table 6.** Putative *Brachypodium distachyon* photoreceptor gene primary transcripts and aliases identified previously and by amino acid sequence comparison with *Arabidopsis thaliana*.

| Gene | *B. distachyon* | *A. thaliana* | Phase (h) |  |  |  |  |  |  |  |
| --- | --- | --- | --- | --- | --- | --- | --- | --- | --- | --- |
|  |  |  | *A. thaliana* |  |  |  | *B. distachyon* |  |  |  |
|  |  |  | LDHC | LDHH | LLHC | LLHH | LDHC | LDHH | LLHC | LLHH |
| CRY1 | Bradi3g46590.1 | At4g08920 | 3 | 0 | 7 | - | 20.7 | 21.3 | - | - |
| CRY2 | Bradi3g49204.4 | At1g04400 | 8 | 15 | 10 | 11 | - | - | 5.9 | 10.1 |
| PHOT1 | Bradi4g45310.2 | At3g45780 | 6 | - | 8 | 7 | - | - | - | - |
| PHOT2 | Bradi5g07360.3 | At5g58140 | 2 | - | 21 | 22 | 21.8 | 20.6* | - | - |
| PHYA | Bradi1g10520.1 | At1g09570 | - | 23 | 10 | 7 | 5.1 | - | 7.1 | - |
| PHYB | Bradi1g64360.1 | At2g18790 | 1 | - | - | 22 | - | - | - | - |
| PHYC | Bradi1g08400.1 | At5g35840 | 8 | - | - | - | - | - | 8.8 | - |
| UVR8 | Bradi5g10580.1 | At5g63860 | 11 | 11 | 13 | 12 | 11.1 | 10.8 | 10.9 | - |

Where *B. distachyon* values are present phase was significant at $p < 0.01$ excluding where \* $p < 0.05$ . – not significant. Source of *A. thaliana* phase calls Michael et al. 2008.

**Supporting Information Table 7.** Wilcoxon rank sum tests comparing period lengths between conditions for circadian transcripts. Benjamini-Hochberg method was used for multiple testing correction. Time course names are as described for Figure 1.

|  | LDHC | LDHH | LLHC |
| --- | --- | --- | --- |
| LDHH | 0.082 | - | - |
| LLHC | $5.9 \times 10^{-6}$ | $7.9 \times 10^{-7}$ | - |
| LLHH | $< 2 \times 10^{-16}$ | $< 2 \times 10^{-16}$ | $< 2 \times 10^{-16}$ |

**Supplemental Table 8.** Wilcoxon rank sum tests comparing relative amplitudes between conditions using Benjamini-Hochberg method for multiple testing correction. Time course names described for Figure 1.

|  | LDHC | LDHH | LLHC |
| --- | --- | --- | --- |
| LDHH | $< 2 \times 10^{-16}$ | - | - |
| LLHC | 0.0086 | $< 2 \times 10^{-16}$ | - |
| LLHH | $< 2 \times 10^{-16}$ | $< 2 \times 10^{-16}$ | $< 2 \times 10^{-16}$ |

**Supporting Information Figure 1.**
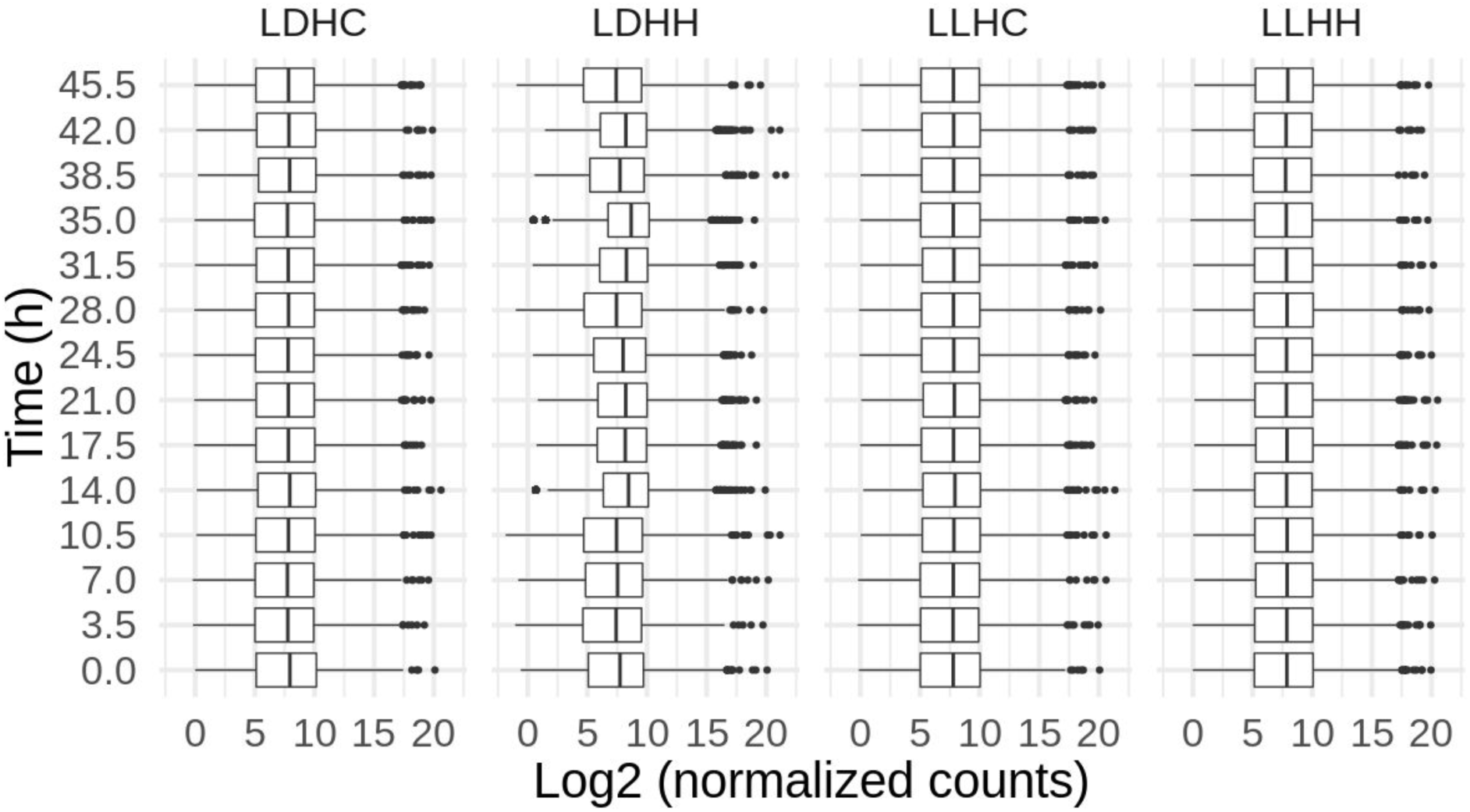
Box and whisker plots showing distribution of normalized sequence read counts per time point across samples. The boxes represent the interquartile range, the median value is depicted by a black line, whisker length is determined as 1.5 times the interquartile range, outliers are represented by separated dots. LDHC, photo- and thermocycles; LDHH, photocycles and constant temperature; LLHC, thermocycles and constant light; LLHH, constant light and warm temperature.

**Supporting Information Figure 2.**
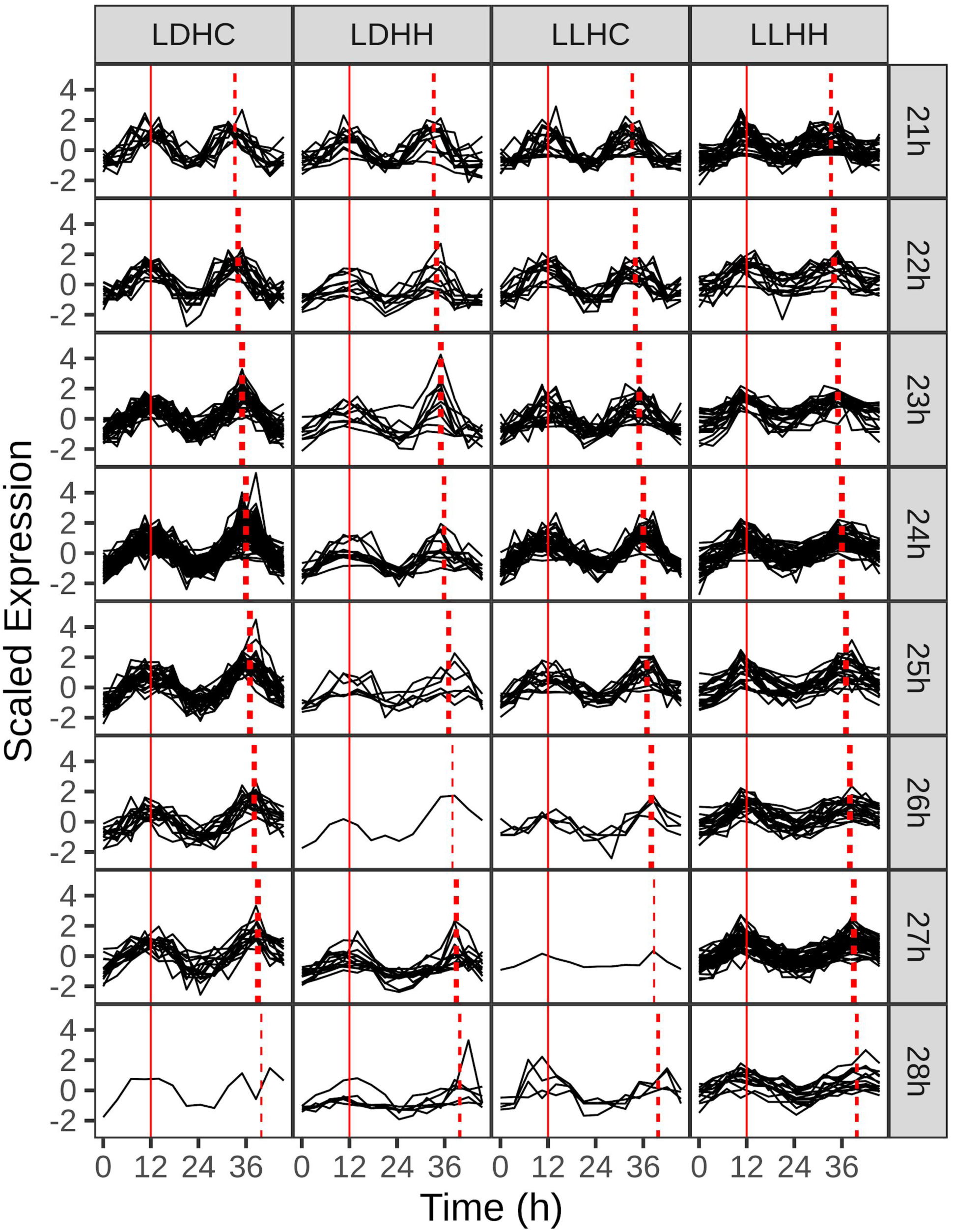
Comparison of period and phase estimates for all cycling transcripts with a phase estimate of approximately 12 h. Transcripts were binned by condition (columns) and period estimate (row). Solid red line depicts rounded phase estimate, dashed red lines depict Metacycle period estimates relative to the phase for all transcripts shown in bin. Time course names as described in Figure 1.

**Supporting Information Figure 3.**
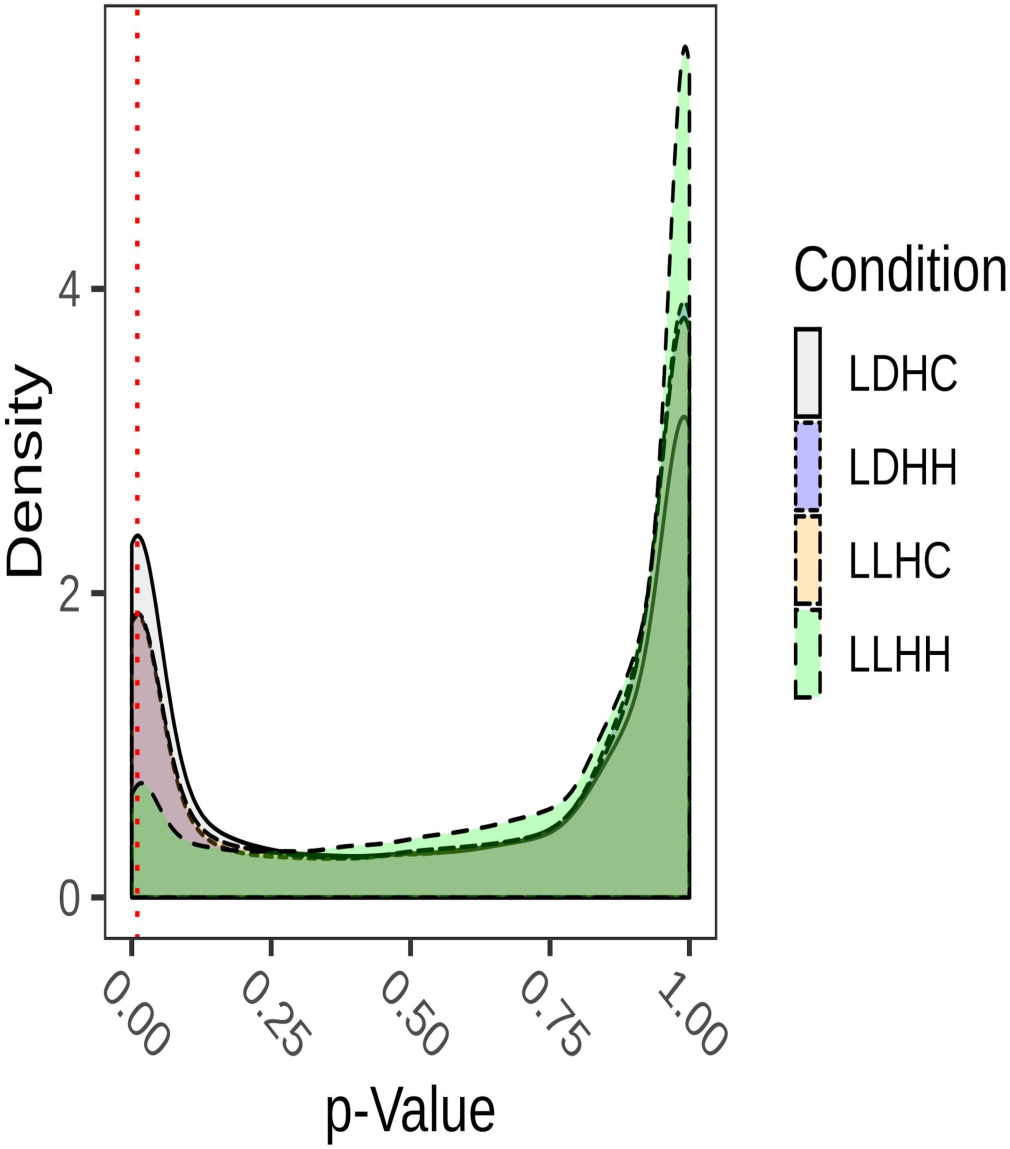
Distribution of the p-values for each transcript in each condition from the MetaCycle test for detecting periodicity. Dotted red vertical line represents cutoff of 0.01. Time course names described in Figure 1.

**Supporting Information Figure 4.**
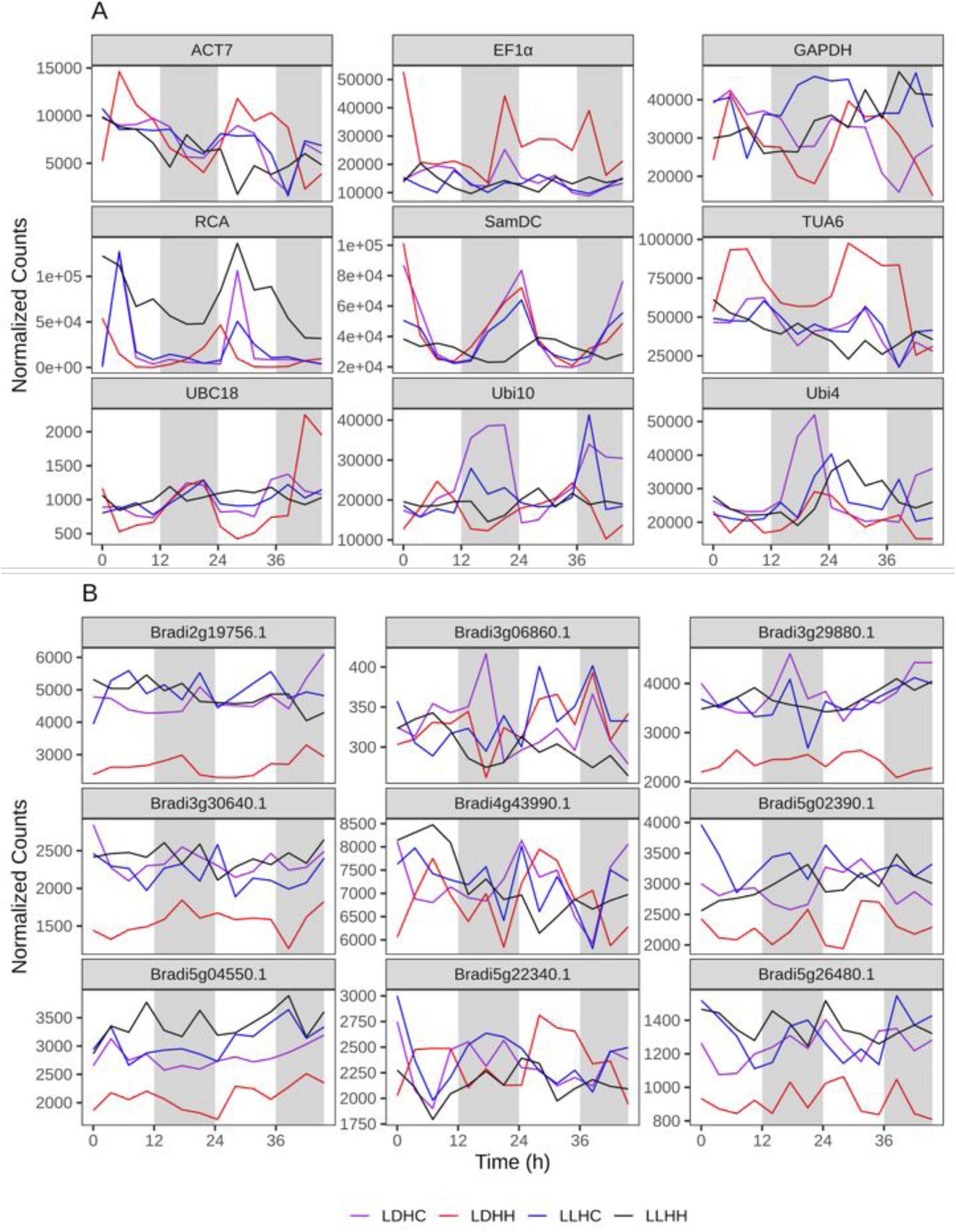
Relative expression of transcripts for use as controls for single-gene PCR quantitation. Relative expression of nine transcripts previously suggested as quantitative reverse transcriptase PCR control genes (A) and transcripts identified as having some of the lowest variance across time courses and across a range of expression levels (B). Time course names are as described in Figure 1.

**Supporting Information Figure 5.**
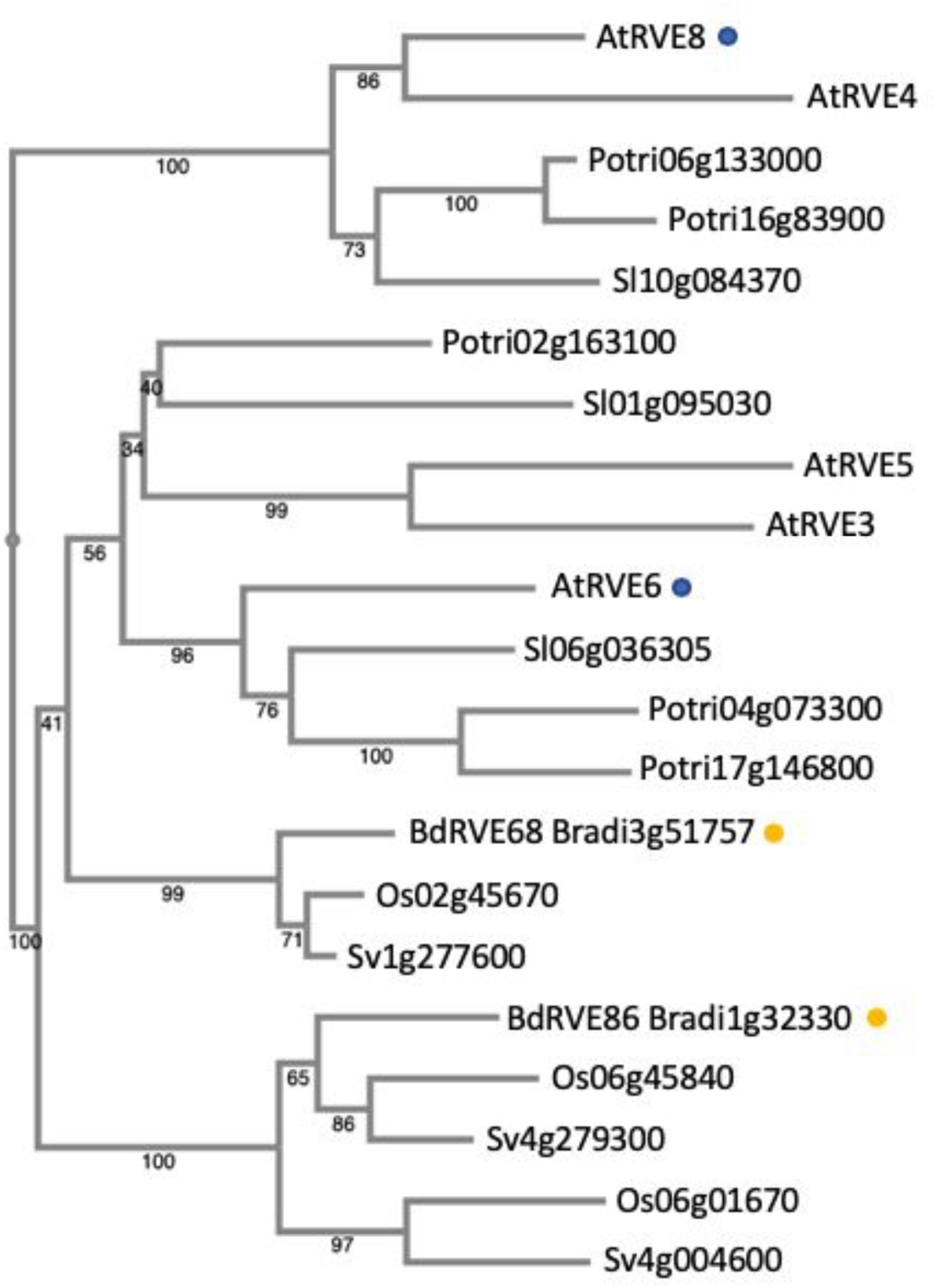
Phylogenetic analysis of *Arabidopsis thaliana REVEILLE* (*RVE*) genes *AtRVE6* and *AtRVE8*, blue dots, and related sequences. Described *Brachypodium distachyon* genes marked by gold dot. The phylogeny was reconstructed using MAFFT (multiple alignment using fast Fourier transform). The numbers labeled on the branches are the posterior probability to support each clade. The branch lengths represent the amino acid substitutions per site. The protein sequences, full locus IDs, and plant species with genome source are provided in Supporting Information Table 2.

**Supporting Information Figure 6.**
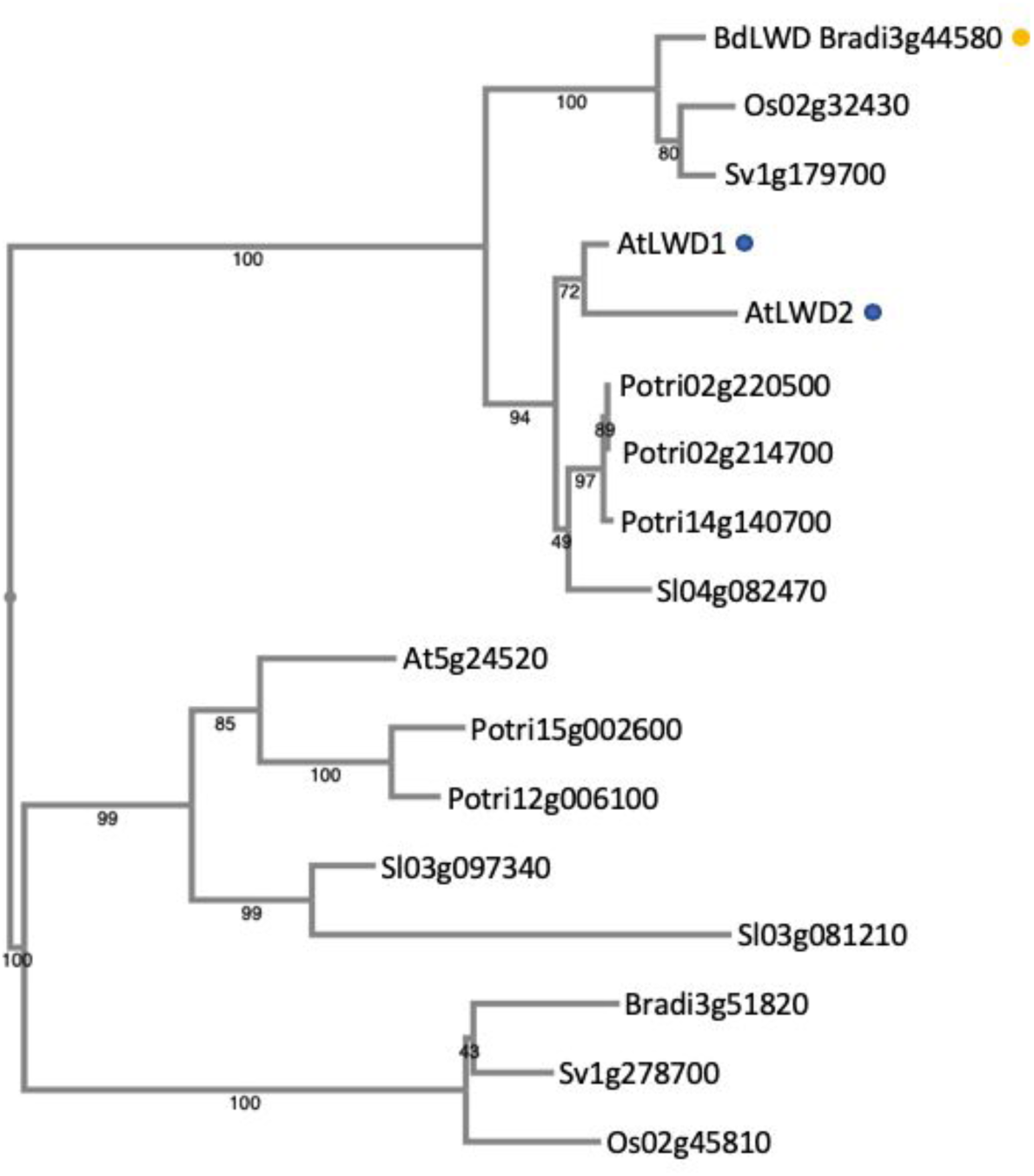
Phylogenetic analysis of *Arabidopsis thaliana LIGHT REGULATED WD* (*LWD*) genes AtLWD1 and AtLWD2, blue dots, and related sequences. Described *Brachypodium distachyon* genes marked by gold dot. The phylogeny was reconstructed using MAFFT (multiple alignment using fast Fourier transform). The numbers labeled on the branches are the posterior probability to support each clade. The branch lengths represent the amino acid substitutions per site. The protein sequences, full locus IDs, and plant species with genome source are provided in Supporting Information Table 2.

**Supporting Information Figure 7.**
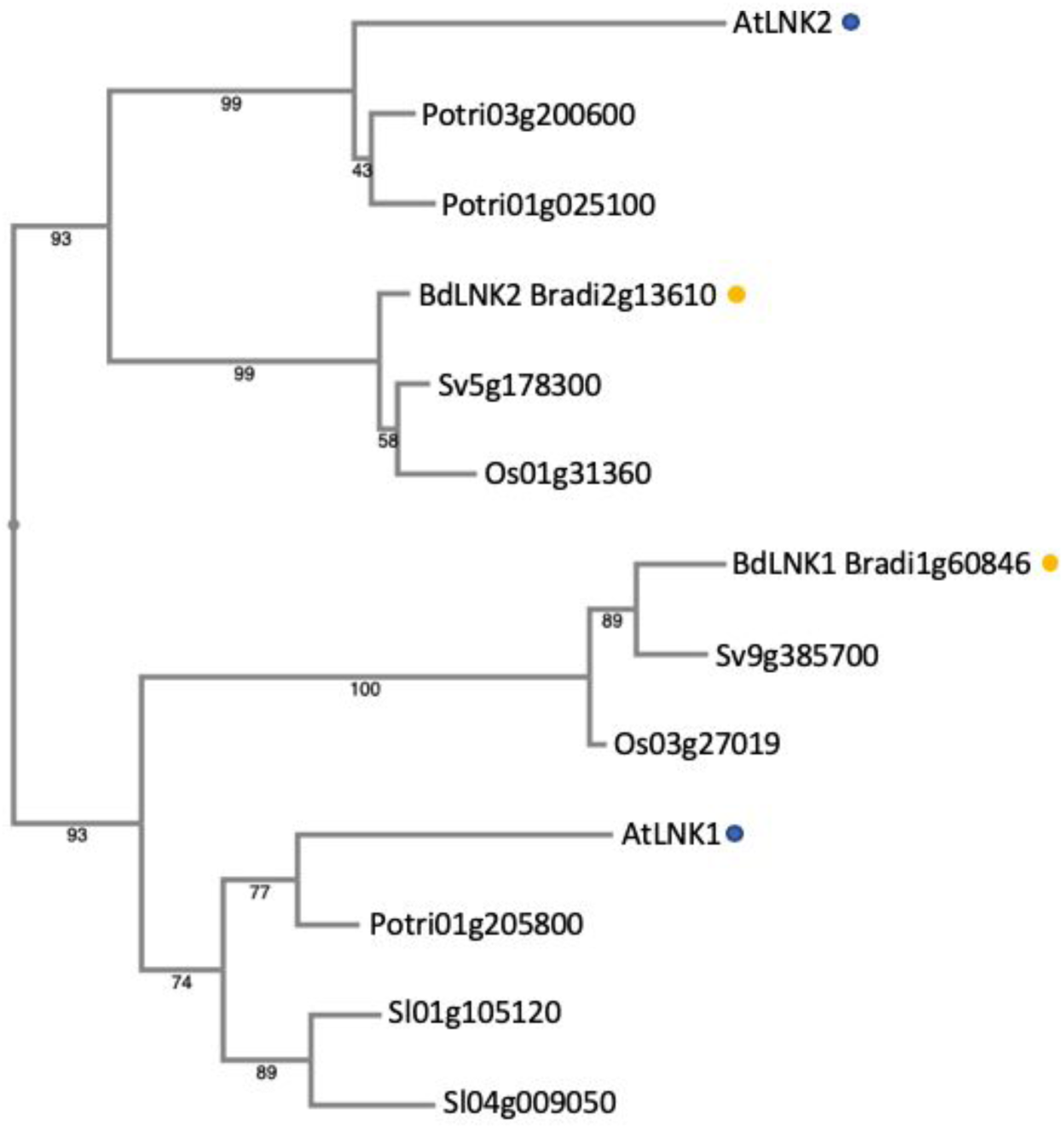
Phylogenetic analysis of *Arabidopsis thaliana NIGHT LIGHT-INDUCIBLE AND CLOCK REGULATED* (*LNK*) genes *AtLNK1* and *AtLNK2*, blue dots, and related sequences. Described *Brachypodium distachyon* genes marked by gold dot. The phylogeny was reconstructed using MAFFT (multiple alignment using fast Fourier transform). The numbers labeled on the branches are the posterior probability to support each clade. The branch lengths represent the amino acid substitutions per site. The protein sequences, full locus IDs, and plant species with genome source are provided in Supporting Information Table 2.

**Supporting Information Figure 8.**
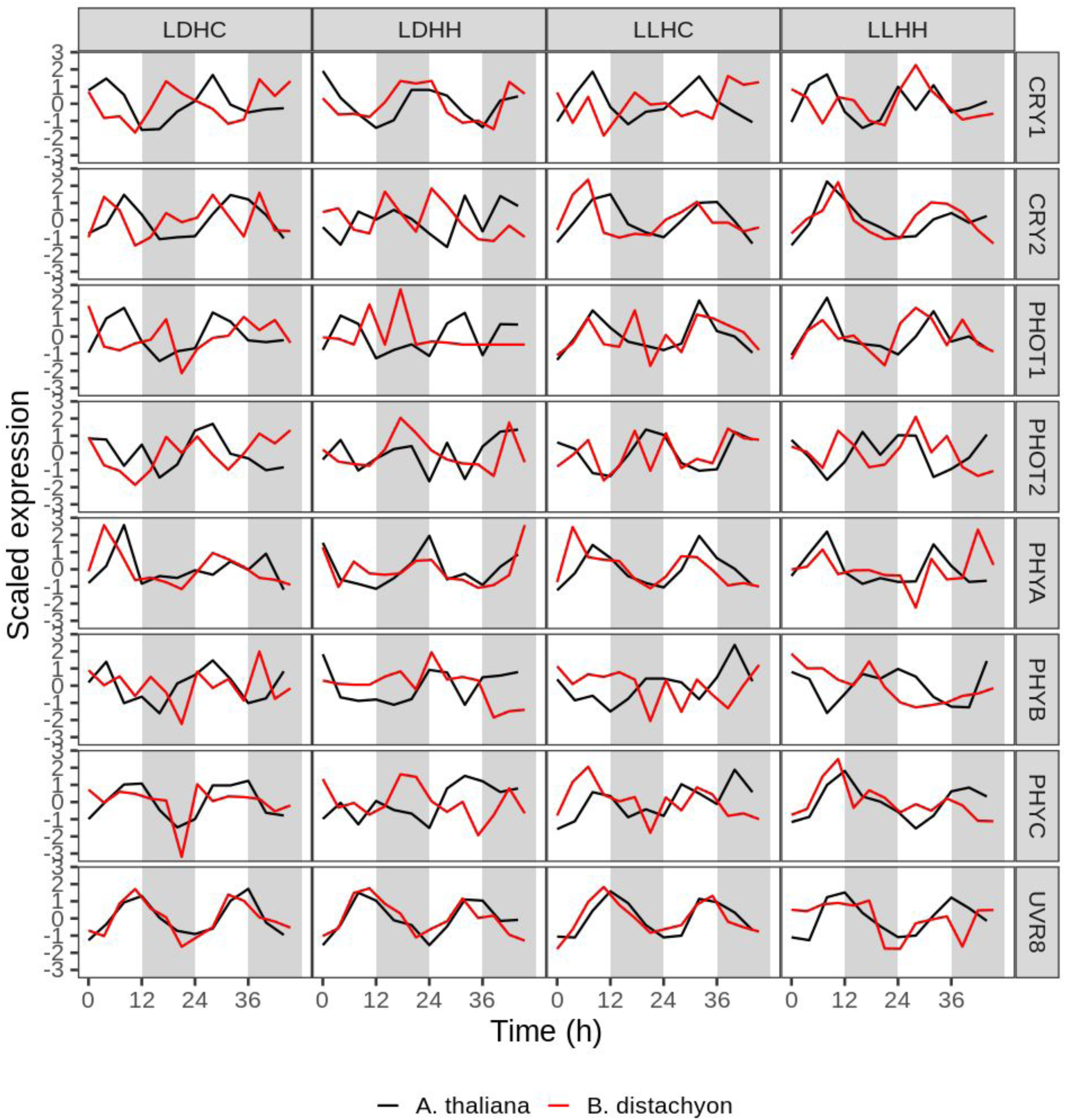
Comparison of *A. thaliana* (black) and *B. distachyon* (red) photoreceptor expression profiles. Condition names at the top of the figure are as described in Figure 1. Gene names are shown to the right of the figure. Gene identification was done through reciprocal best hit.

**Supporting Information Figure 9.**
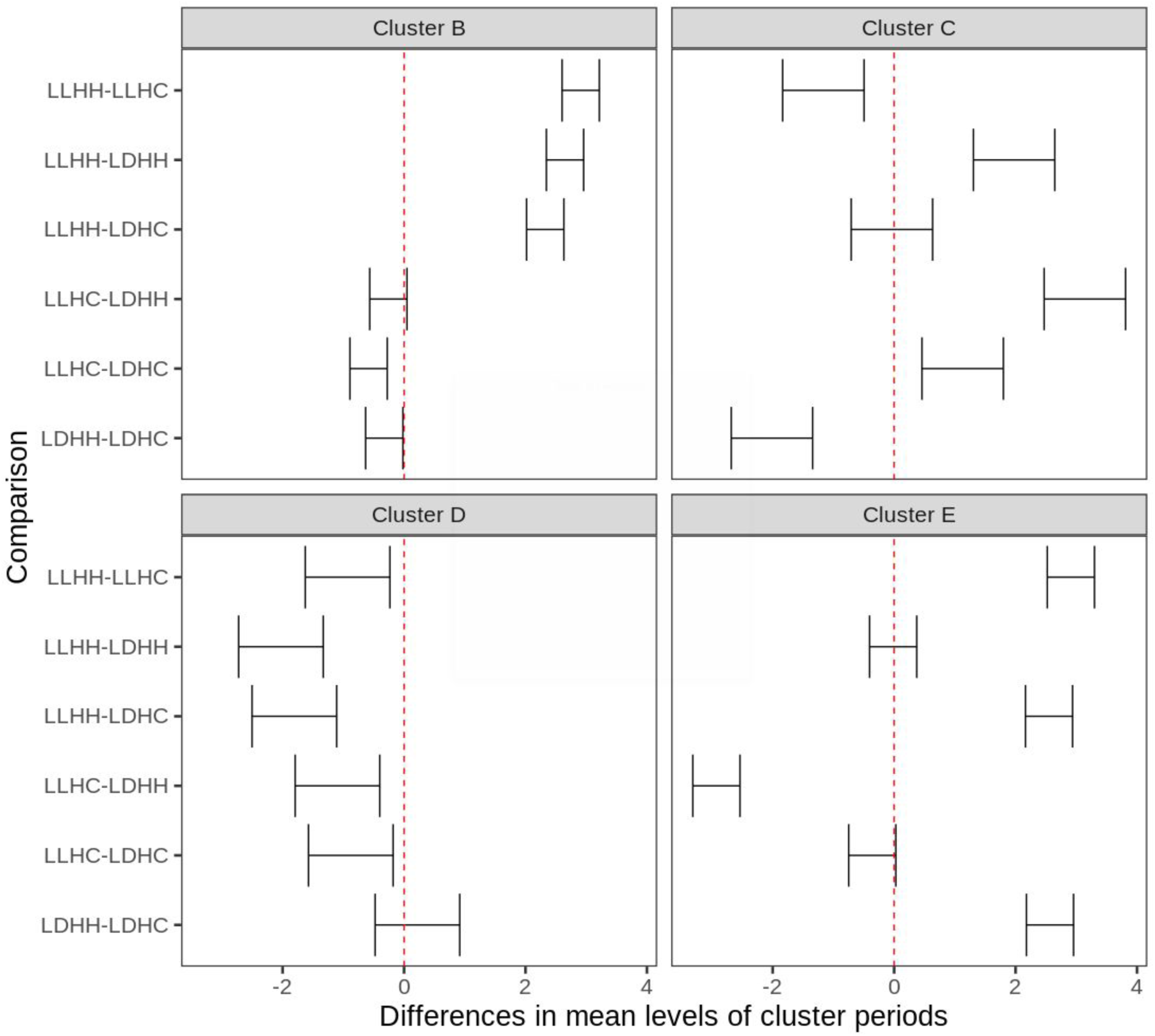
95% family-wise confidence level of mean cluster periods from post-hoc Tukey’s honest significant differences test. Ranges overlapping the red dashed line (0) indicate non-significant differences in means. Cluster names refer to panel position in Figure 4. Condition names are as described in Figure 1.

**Supporting Information Figure 10.**
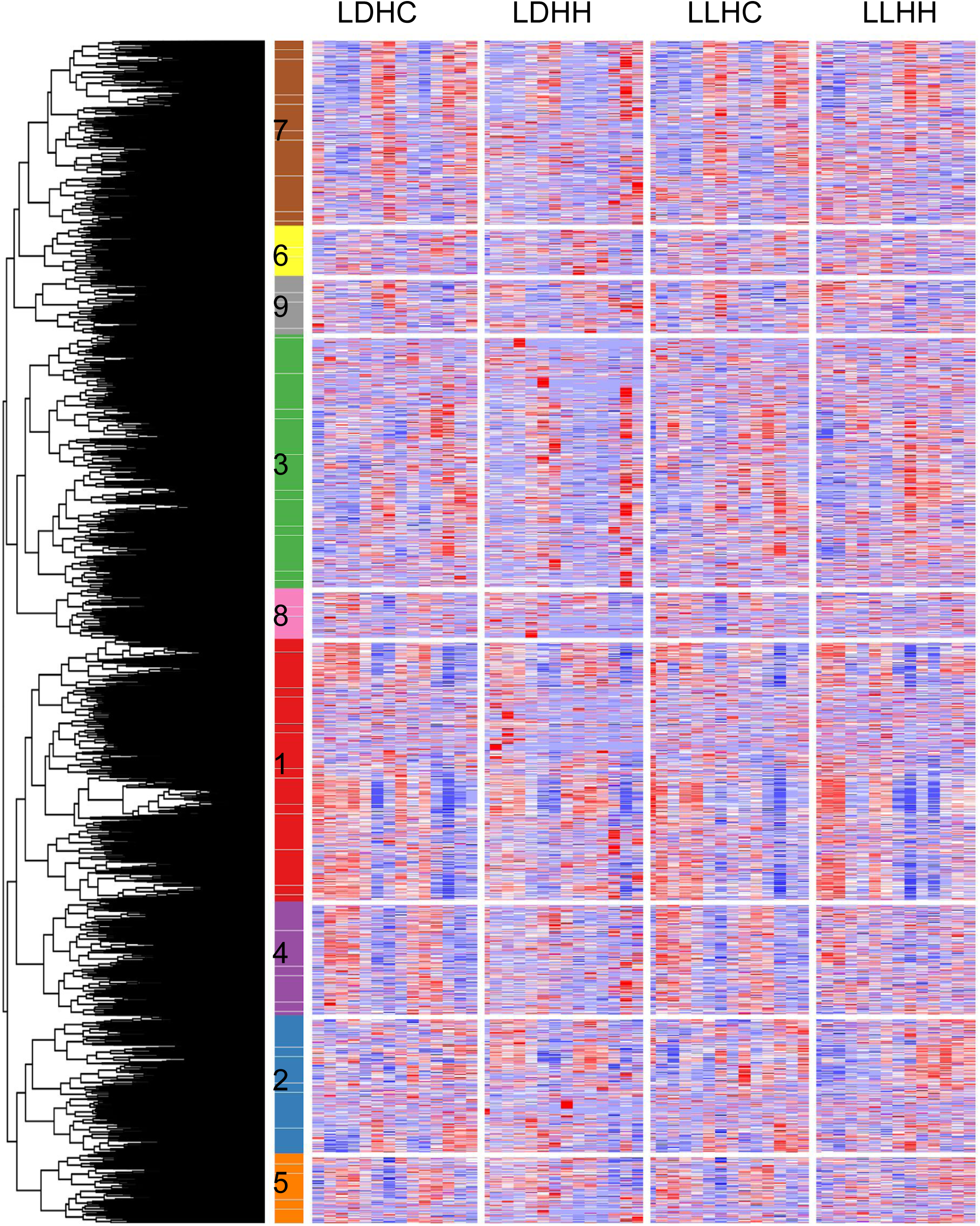
Hierarchical clustering of transcripts shows conservation of expression under thermocycles. All transcripts expressed in each of the four conditions (n=39,048) were hierarchically clustered using Pearson correlation with complete linkage. The nine distinct cluster groups were decided using the elbow method. Red shading is indicative of greater expression and blue shading implies decreased expression. Clusters are denoted by a colored vertical bar and numbered for future reference. Time course names described for Figure 1.

**Supporting Information Figure 11.**
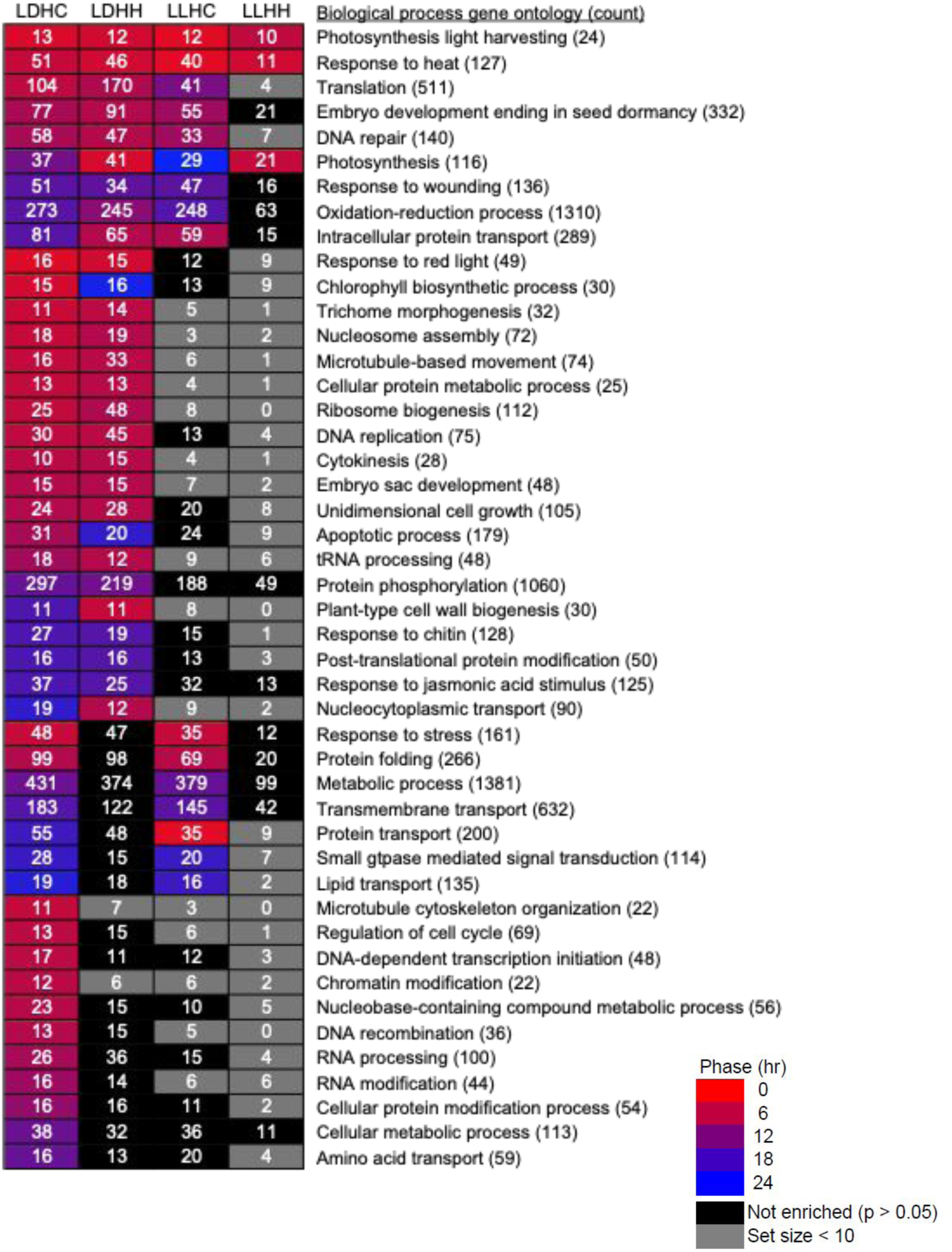
Time-of-day expression of biological process gene ontology terms were often uniquely regulated by time course conditions. Transcripts were grouped based on phase and tested for biological process gene ontology terms using the indertifiers of the reciprocally most similar *Arabidopsis thaliana* homolog. Values are the count of rhythmic transcripts in each phase bin and color gradient indicates the average phase of significantly enriched sets (*P* < 0.05). Sets containing fewer than 10 rhythmic genes were due to the small sample size (grey). Time course names described for Figure 1.

**Supporting Information Figure 12.**
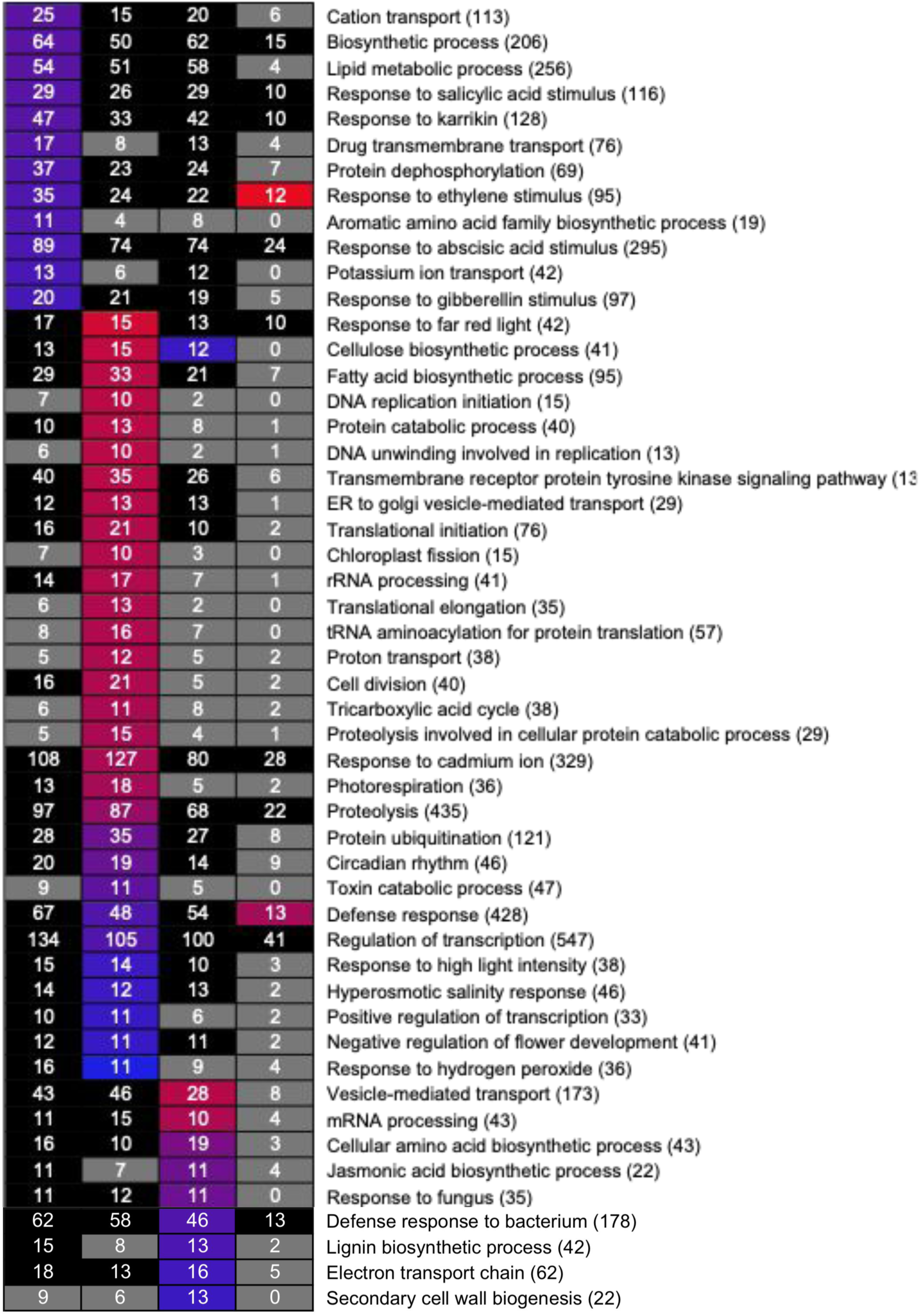
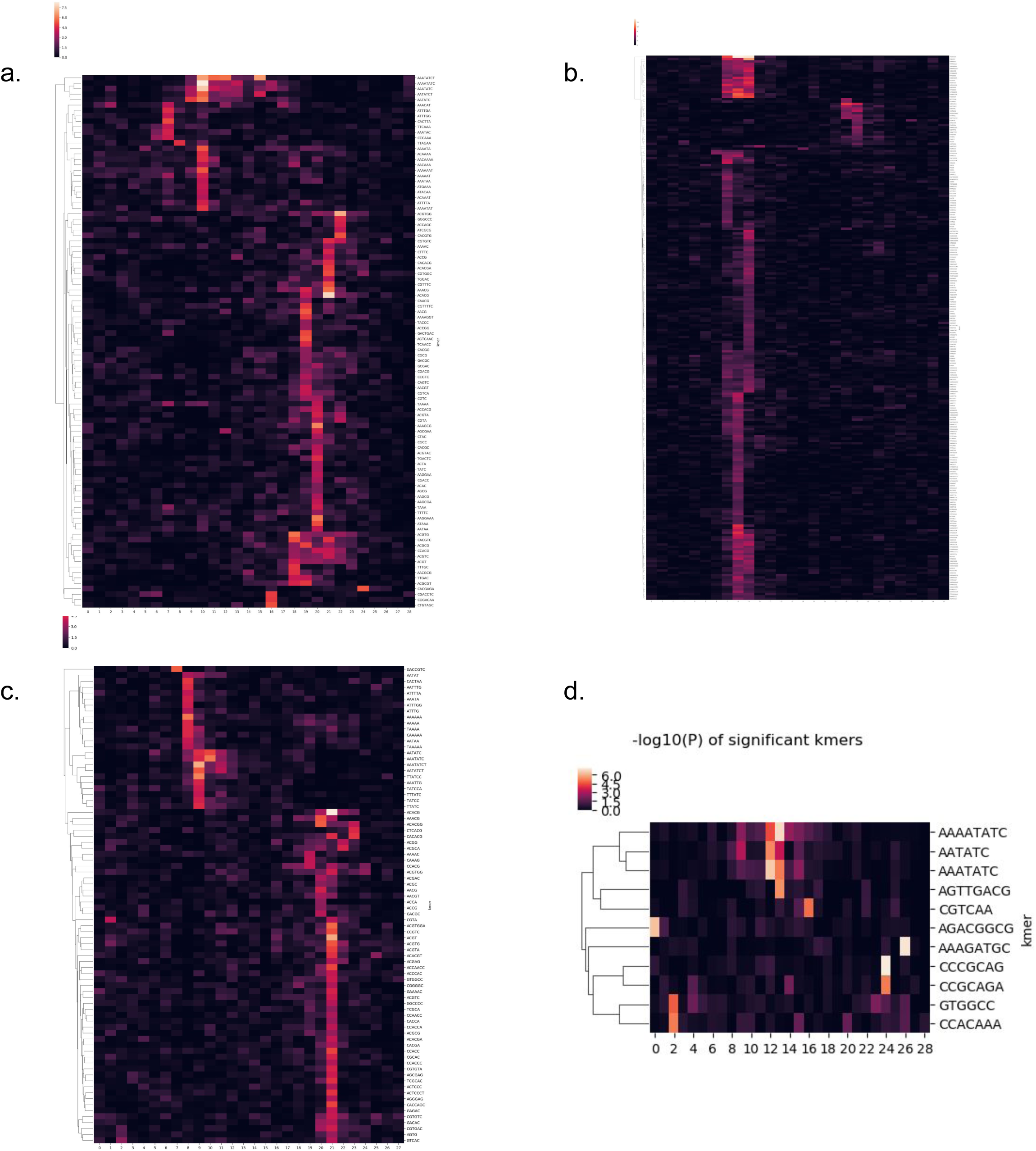
Heat map of significant 3-8 bp elements across diurnal and circadian conditions. Significant 3-8 mers under (a) LDHC, (b) LDHH, (c) LLHC and (d) LLHH were clustered based on the time of their overrepresentation Z score. Dendrogram on the left and the 3-8 bp element sequence on the right.

**Supporting Information Figure 13.**
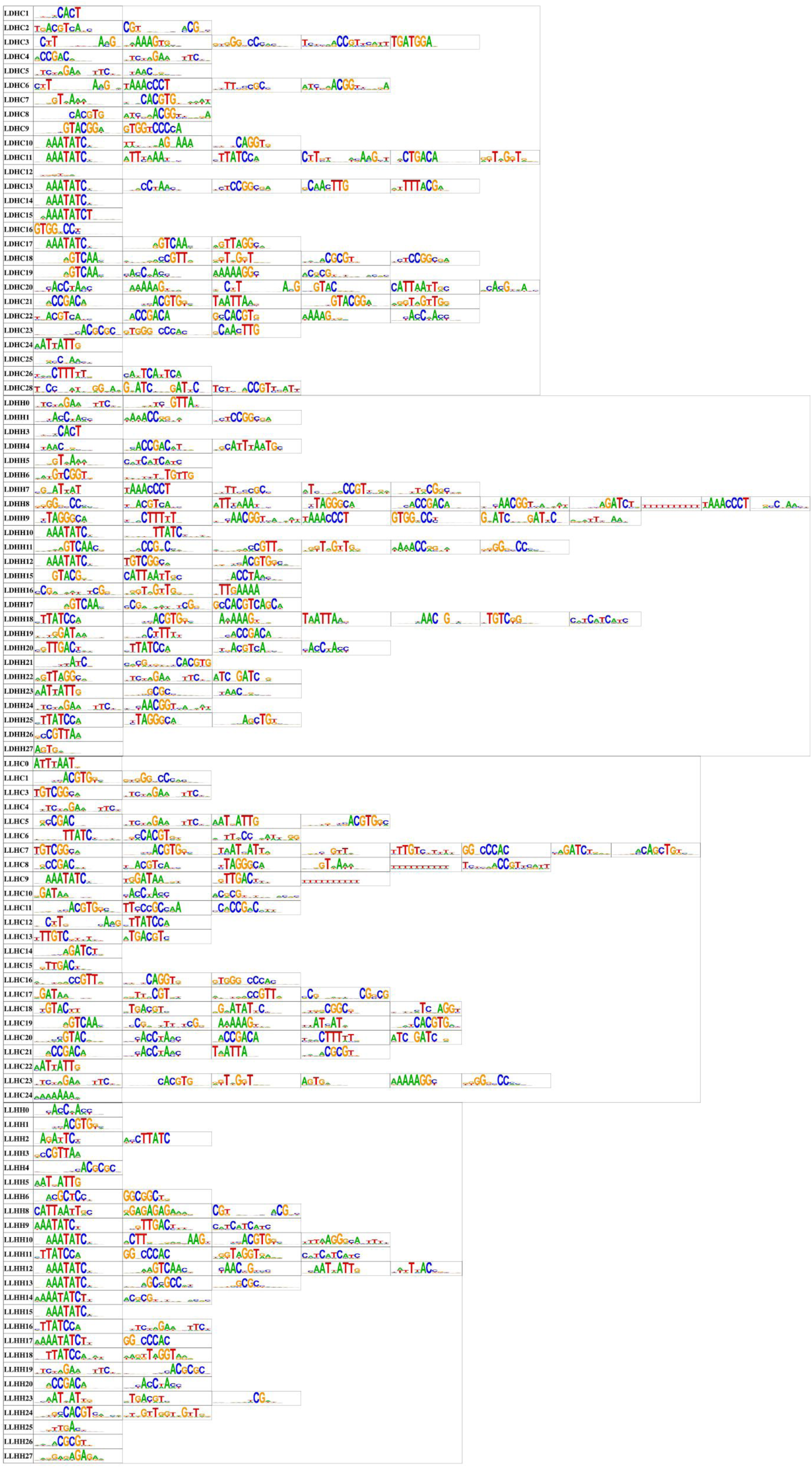
Complete list of *cis*-regulatory sequences most overrepresented at each hour in each time course. Each nucleotide sequence is a position probability matrix motif derived from DNA-affinity purification sequencing. The height of the letters at each position is proportional to the probability of a nucleotides. Time course names described for Figure 1.

**Supporting Information Figure 14.**
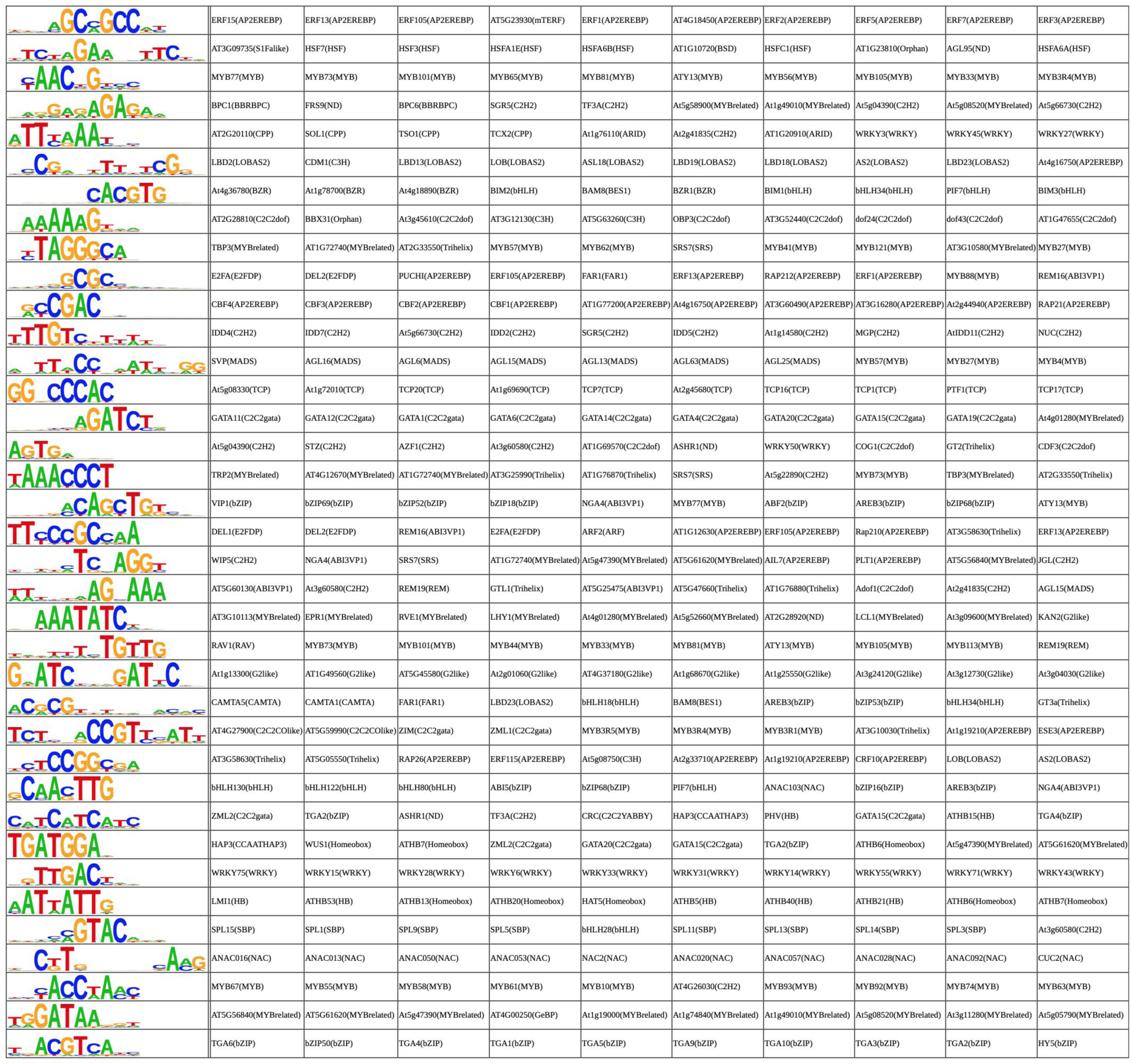
Identification of transcription factor proteins and the greatest likelihood matching DAP-seq sequence logos that they bind. Each nucleotide sequence is a position probability matrix motif derived from DNA-affinity purification sequencing. Time course names described for Figure 1.

